# Herbaceous production lost to tree encroachment in United States rangelands

**DOI:** 10.1101/2021.04.02.438282

**Authors:** Scott L. Morford, Brady W. Allred, Dirac Twidwell, Matthew O. Jones, Jeremy D. Maestas, Caleb P. Roberts, David E. Naugle

## Abstract

1. Rangelands of the United States provide ecosystem services that benefit society and rural economies. Native tree encroachment is often overlooked as a primary threat to rangelands due to the slow pace of tree cover expansion and the positive public perception of trees. Still, tree encroachment fragments these landscapes and reduces herbaceous production, thereby threatening habitat quality for grassland wildlife and the economic sustainability of animal agriculture.
2. Recent innovations in satellite remote sensing permit the tracking of tree encroachment and the corresponding impact on herbaceous production. We analyzed tree cover change and herbaceous production across the western United States from 1990 to 2019.
3. We show that tree encroachment is widespread in U.S. rangelands; absolute tree cover has increased by 50% (77,323 km^2^) over 30 years, with more than 25% (684,852 km^2^) of U.S. rangeland area experiencing tree cover expansion. Since 1990, 302 ± 30 Tg of herbaceous biomass have been lost. Accounting for variability in livestock biomass utilization and forage value reveals that this lost production is valued at between $4.1 - $5.6 billion U.S. dollars.
4. Synthesis and applications: The magnitude of impact of tree encroachment on rangeland loss is similar to conversion to cropland, another well-known and primary mechanism of rangeland loss in the U.S. Prioritizing conservation efforts to prevent tree encroachment can bolster ecosystem and economic sustainability, particularly among privately-owned lands threatened by land-use conversion.

## 1. Introduction

Native trees are invading global rangeland biomes (Nackley et al., 2017). Fire suppression, livestock overgrazing, nutrient pollution, and increasing CO_2_ emissions contribute to extensive tree encroachment in rangelands (here defined as grasslands, shrublands, and open woodlands) (Asner et al., 2004). This encroachment exacerbates already substantial losses of rangelands to cropland conversion and threatens global rangelands’ resilience and conservation potential (Fargione et al., 2018; Hoekstra et al., 2004; Kremen & Merenlender, 2018; Veldman et al., 2015).

Tree encroachment modifies the structure and function of rangeland ecosystems, thereby impacting a host of ecosystem services. Water storage and supply, wildlife and livestock habitat quality, biodiversity preservation, and climate regulation are all modulated by tree cover in rangelands (Archer and Predick, 2014; Bardgett et al., 2021). Importantly, tree cover is a key regulator of herbaceous production (the combined production of grasses and forbs; i.e., forage) upon which both wildlife and livestock depend. Substantial effort has been invested in disentangling the complex relationship between herbaceous production and tree cover (Anadon et al., 2014; Scholes, 2003). Yet, critical knowledge gaps still exist for how herbaceous production responds to tree encroachment at large scales.

Identifying where tree encroachment impacts ecosystem services such as carbon (C) storage and productivity has traditionally relied on historical image analysis, paired site investigations, and evaluations of the pollen record (Archer et al., 1988; Lubetkin et al., 2017). These methods have documented substantial tree canopy expansion in Africa, the Americas, Asia, and Australia. Unfortunately, critical knowledge on patterns, pace, and magnitude of tree encroachment to rangelands remains absent at large scales.

While operational remote sensing products are applied globally to track forest cover changes, these analyses typically lack the sensitivity to track tree cover gains in rangelands or omit analysis of tree cover in rangelands altogether (Hansen et al., 2013; Rigge et al., 2020).

Recent technological advances in remote sensing collectively suggest large-scale tree cover expansion in United States (U.S.) rangelands (Allred et al., 2021; Filippelli et al., 2020; Jones et al., 2020), consistent with decades of empirical observations. Related innovations in tracking annual herbaceous production now allow for an integrated spatiotemporal analysis of production and tree cover (Jones et al., 2021)

Two-thirds of western U.S. rangelands are under private ownership, and conservation of these lands requires addressing ecological and economic sustainability linkages. Declining ranch income hastens the conversion of intact grasslands to row-crop agriculture, fossil-fuel energy production, and dispersed housing developments (Allred et al., 2015; Lark, 2020). For example, row-crop conversion has accelerated in recent years due to commodity market fluctuations and declining economic returns from livestock production. Tree encroachment can exacerbate declines in ranching profitability as expanding woodlands out-compete and displace grasses and forbs (Anadon et al., 2014; Fuhlendorf et al., 2008). Revealing the economic cost of tree encroachment on forage production could help motivate livestock producers to implement proactive, conservation-focused tree management in rangelands.

Here we investigate tree encroachment and its impacts to herbaceous production across primary U.S. rangelands. We estimate the economic value of lost production to help inform sustainable agricultural practices, the preservation of intact grasslands (Scholtz & Twidwell, 2022), and the protection of grassland bird habitats threatened by tree encroachment (Baruch-Mordo et al., 2013; Herse et al., 2018; Rosenberg et al., 2019). Specifically, our analysis quantifies the difference between attainable and actual herbaceous production as a function of tree cover expansion, a difference termed yield gap that is commonly used in agriculture to investigate production losses (van Ittersum et al., 2013). Yield gap estimates were combined with grazing rental rates from the U.S. Department of Agriculture (USDA) to estimate foregone revenue to livestock producers from tree encroachment.

## 2. Methods

We calculated tree cover change and herbaceous production lost to tree encroachment using data from the Rangeland Analysis Platform version 2.0 (RAP-V2, Allred et al., 2021). RAP-V2 geography spans 17 western states (**Fig. 1**) and includes the entirety of intact semi-arid and arid rangelands in the U.S. (Reeves & Mitchell, 2011, Scholtz & Twidwell, 2022). Our analysis incorporated data from 1990 through 2019. We used Landfire Biophysical Settings (BPS) version 1.4.0, to identify where rangelands were dominant prior to Euro-American settlement and to exclude primary forest areas from our calculations. Our analysis also excludes contemporary lands that were mapped as croplands, urban/built, or riparian. This analysis included 2,783,955 km^2^ of historical rangelands.

**Figure 1.**
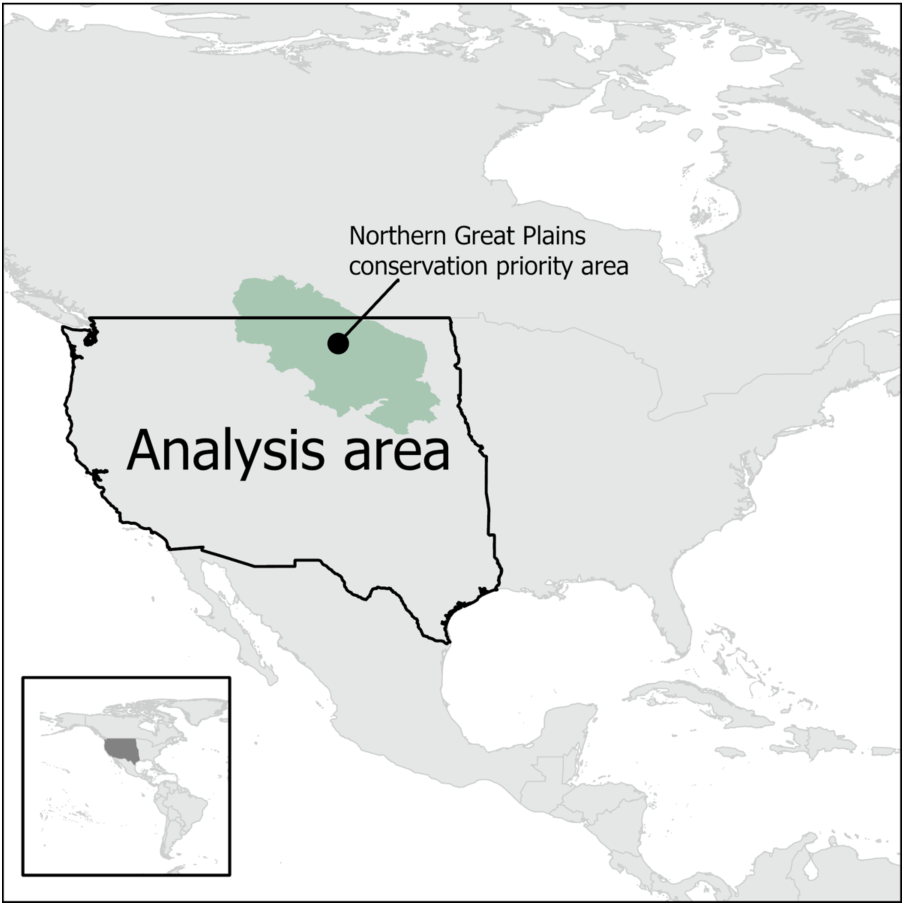
Yield gap analysis area includes the western half of the United States of America, covering approximately 2.8 million km^2^ of rangelands.

The multi-part modeling framework is presented below in four sections. Tree cover and herbaceous production modeling are presented first. These data were used as inputs to the yield gap modeling presented in section 2.3. Then, section 2.4 describes how the yield gap biomass estimates were combined with economic data to produce a monetary estimate of lost production.

### 2.1 Tree Cover Modeling and Analysis

In RAP-V2, fractional tree cover was calculated using a convolutional neural network. The network used Landsat surface reflectance (bands 1-5,7 for Landsat 5 TM and 7 ETM+; bands 2-7 for Landsat 8 OLI), normalized difference vegetation index (NDVI), and normalized burn ratio two (NBR2) for prediction. The model was trained using data from 52,012 vegetation field plots collected by Federal natural resource agencies. Model performance was evaluated against 5,780 field plots; metrics evaluated include mean absolute error (MAE = 2.8%), root mean square error (6.8%), residual standard error (5.9%) and coefficient of determination (r2 = 0.65).

To improve the detection of real and significant tree cover change, we further processed the tree cover data using the LandTrendr (LT) segmentation algorithm (Kennedy et al., 2010) in Google Earth Engine (GEE). The LT parameters used in this analysis are summarized in **Supplemental Table 1**. We applied two rules to remove spurious tree cover trends in our analysis. First, to tally tree cover change, the difference in tree cover needed to exceed the model MAE (2.8%). Second, we performed a Welch T-test (n = 4, two-tailed, p-value = 0.01) on each pixel segment using the RAP-V2 uncertainty data. For tree cover change to be tallied in a pixel, it had to have one or more segments where the t-value < p-value.

*Tree-free rangelands* in this analysis were defined functionally at the pixel level (30m by 30m) as rangelands where tree cover was less than 4%, corresponding to tree cover where sizable impacts to ecosystem function are observed (Baruch-Mordo et al., 2013). Tree-free rangelands were further categorized as either *intact* or *vulnerable. Vulnerable tree-free rangelands* were classified as *tree-free rangeland* pixels where tree cover was greater than 2.8% (the tree cover MAE) in pixels within a 200-meter radius. In our analysis we evaluate both the change in total tree-free rangelands on a per-pixel basis and change in intact/vulnerable tree-free rangelands based on the neighborhood analysis.

### 2.2 Herbaceous Production Calculations

Pixel-level estimates for herbaceous production were calculated using the Landsat implementation of the MOD17 Net Primary Productivity algorithm and calibrated for aboveground herbaceous production (Jones et al., 2021; Robinson et al., 2019). The algorithm incorporated land surface reflectance, land surface cover from RAP-V2, and meteorology data. Production calculations from this method show good agreement with independent estimates of production from remote sensing and field-based compilations (r^2^ values of 0.79 and 0.82, respectively), but tend to under-estimate production in highly productive environments (See Figure 2 in Jones et al., 2021 and supplemental text). In the modeling for this analysis, herbaceous production estimates from the MOD17 algorithm are referred to as *Observed Production*.

**Figure 2.**
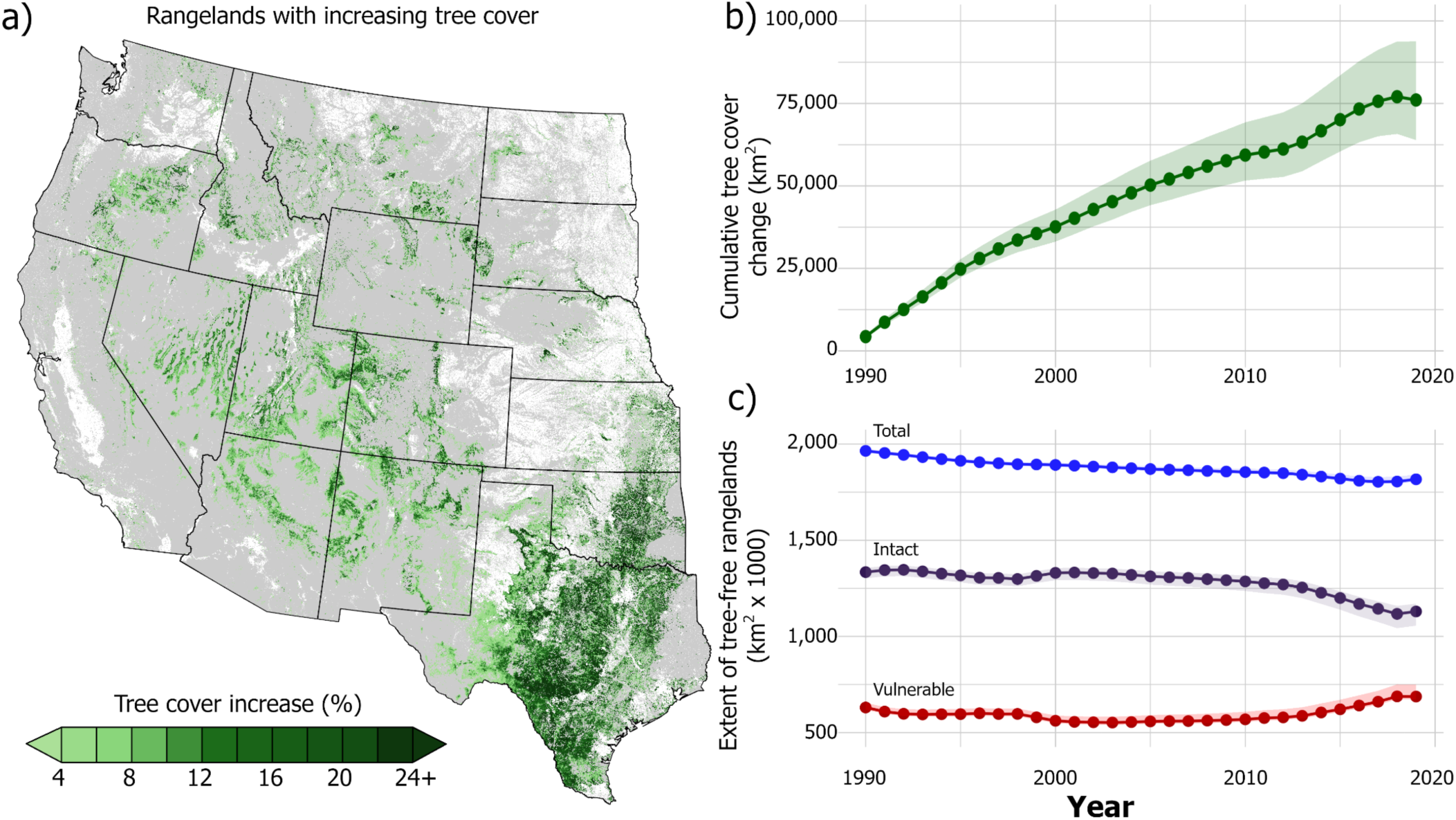
Woody encroachment in western U.S. rangelands (1990 - 2019). a) Net tree canopy expansion in rangelands shown in green; converted agricultural lands and built environment shown in white. b) Cumulative net tree cover expansion totaled 77,323 km^2^ over 30 years; error bands represent the cumulative 95% prediction interval. c) Woodland extensification has resulted in an 8% decrease (147,700 km^2^) of tree-free rangelands in the past 30 years. Intact tree-free rangelands (lands without a tree seed source within 200 meters) have declined more rapidly than the total, showing that tree-free rangelands are increasingly vulnerable to woodland conversion due to the prevalence of nearby tree seed sources.

### 2.3 Yield Gap Modeling

Next, we combined the tree cover and production data presented in sections 2.1 and 2.2 with other biophysical variables to calculate herbaceous production lost to increased tree cover. We used XGBoost (Chen & Guestrin, 2016), an ensemble decision tree supervised learning algorithm, to model herbaceous production as a function of the variables presented in **Table 1**.

**Table 1.**
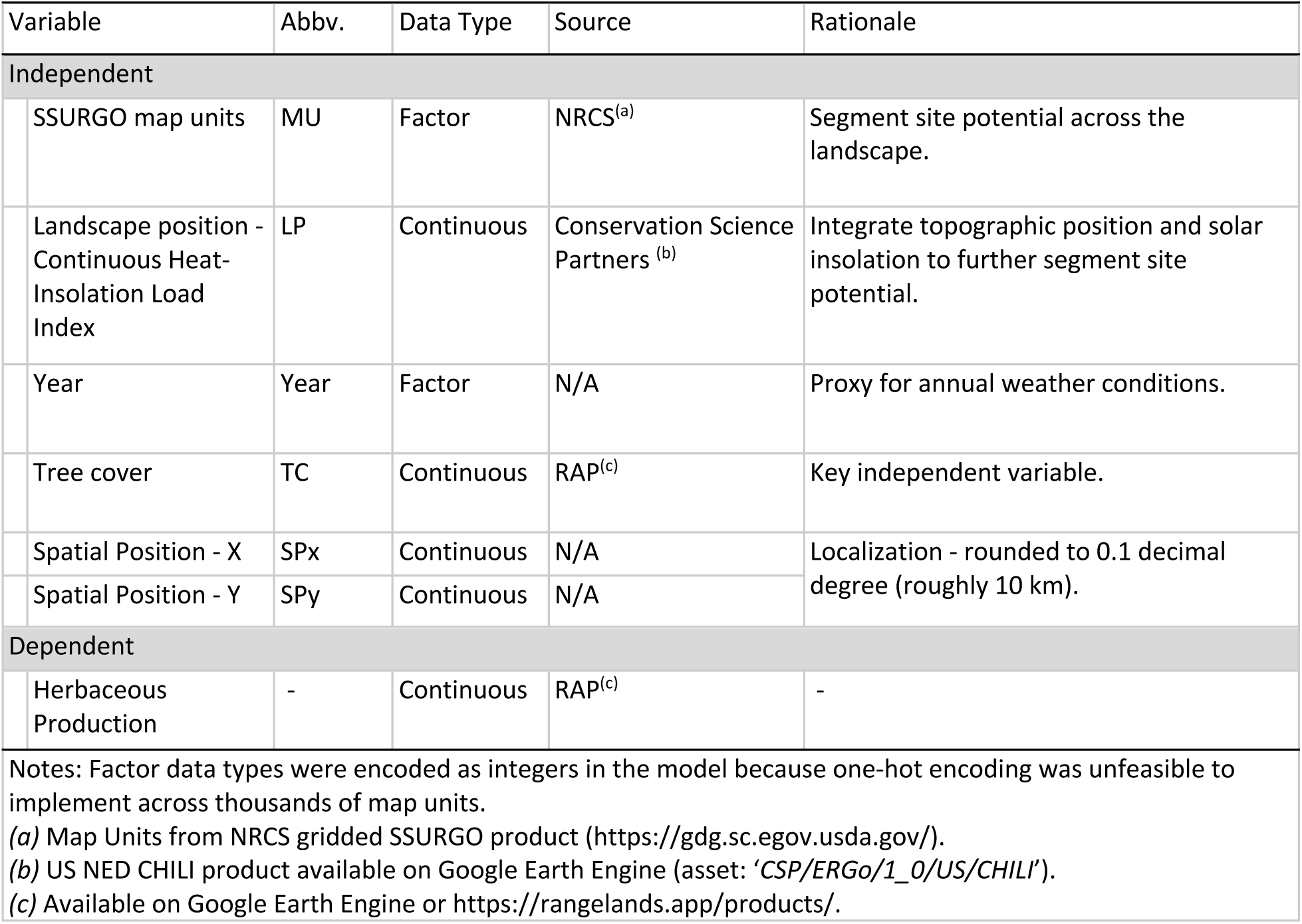
Yield gap modeling variables

| Table 1. Yield gap modeling variables |  |  |  |  |
| --- | --- | --- | --- | --- |
| Variable | Abbv. | Data Type | Source | Rationale |
| Independent |  |  |  |  |
| SSURGO map units | MU | Factor | NRCS<sup>(a)</sup> | Segment site potential across the landscape. |
| Landscape position - Continuous Heat-Insolation Load Index | LP | Continuous | Conservation Science Partners <sup>(b)</sup> | Integrate topographic position and solar insolation to further segment site potential. |
| Year | Year | Factor | N/A | Proxy for annual weather conditions. |
| Tree cover | TC | Continuous | RAP<sup>(c)</sup> | Key independent variable. |
| Spatial Position - X | SPx | Continuous | N/A | Localization - rounded to 0.1 decimal degree (roughly 10 km). |
| Spatial Position - Y | SPy | Continuous | N/A |  |
| Dependent |  |  |  |  |
| Herbaceous Production | - | Continuous | RAP<sup>(c)</sup> | - |
| Notes: Factor data types were encoded as integers in the model because one-hot encoding was unfeasible to implement across thousands of map units. |  |  |  |  |
| (a) Map Units from NRCS gridded SSURGO product (https://gdg.sc.egov.usda.gov/). |  |  |  |  |
| (b) US NED CHILI product available on Google Earth Engine (asset: 'CSP/ERGo/1_0/US/CHILI'). |  |  |  |  |
| (c) Available on Google Earth Engine or https://rangelands.app/products/. |  |  |  |  |

The XGBoost algorithm made two biomass predictions per pixel to estimate yield gap: *Estimated Production* and *Achievable Production. Estimated Production* represents the model’s best estimate of actual biomass production; *Achievable Production* represents potential production, assuming tree cover remains constant over their 30-year modeling domain. The difference between these two production estimates represents the yield gap.

More formally, our analysis applies the following framework to model herbaceous production as a function of tree cover and biophysical variables (**equation 1**).

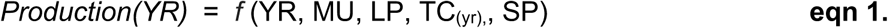

In our framework, *YR* represents year, MU = soil map unit, LP = landscape position, TC_(yr)_ = tree cover for year YR, SP = spatial position/coordinates. Factor variables were encoded as integers during training and inference. Model variables, source data, and a rationale for their inclusion are presented in **Table 1**.

#### Model parameterization and training

We split our analysis area into 53 spatial units using Level-3 EcoRegions from the U.S. Environmental Protection Agency. The training data size for each unit was limited to 71,582,780 records due to computational limitations; the remaining training data were used as hold-out (test) data. The models were trained with 3.8 billion records randomly sampled from our dataset; the remaining 5.5 billion records were used for validation and evaluation. The optimization (loss) function was mean squared error with L2 regularization; our ensemble used 60 trees with a maximum tree depth of 15 **(Supplemental Table 2**). Evaluation of the training metrics showed that holdout MAE values closely matched training MAE values (r^2^ > 0.99), suggesting the model did not overfit the data (**Supplemental Fig. 1)**.

#### Inference

XGBoost generated two predictions: *Estimated Production* and *Achievable Production* ***(equations 2 and 3)***, assuming that tree cover remained constant at 1990 levels.

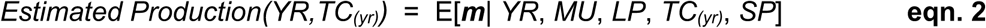

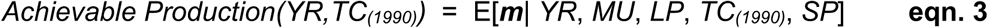

The yield gap attributable to tree cover expansion since 1990 is calculated by subtracting *Estimated Production* from *Achievable Production* for each pixel-year combination (***equation 4***).

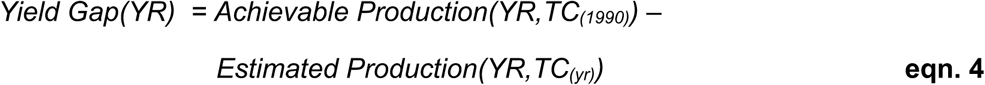

*Prediction Error* for each pixel was calculated as the absolute difference between *Observed Production and Estimated Production* ***(equation 5)***. *Model Error* was represented as the MAE **(equation 6)**, aggregated by year and map unit to provide some localization; *n* is the number of observations for each MU.

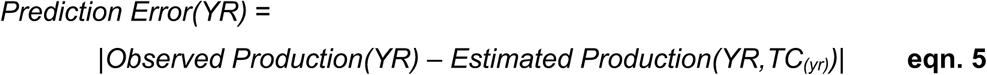

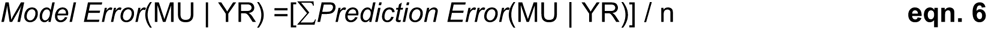

For this analysis, the yield gap was tallied when *Yield Gap* > *Model Error* for 3 or more consecutive years.

### 2.4 Monetary calculations

We calculated the dollar value of herbaceous production lost to tree encroachment by combining yield gap calculated in section 2.3 with estimates with annual state-level grazing rental rates from the USDA National Agricultural Statistics Service (NASS, U.S. Department of Agriculture, 2020). Producer-to-producer rental rates are the most direct way to estimate the market value of herbaceous production. In the U.S., rental contracts are most commonly negotiated using Animal Unit Month (AUM) lease rates. AUM is a measure of forage use by livestock. The value of herbaceous production was calculated from AUM rental rates using **equation 7**.

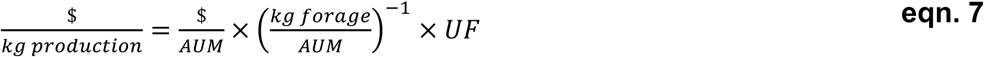

Dollars per Animal Unit Month ($/AUM) represents the grazing rental rate; *kg forage / AUM* represents the mass of consumed forage per AUM and is assumed to equal 344.73 kg forage/AUM. UF is the utilization factor and represents the fraction of total herbaceous production consumed as forage by livestock. The UF is required to convert between total herbaceous production (what is grown) and forage (what is utilized).

The UF is difficult to estimate because it can vary in space and time. Literature based estimates suggest that UF is equal to 0.43 under moderate grazing regimes (Holechek et al., 2010), which generally conforms to the use-half leave-half principle prescribed in U.S. grazing systems. Since 2008, NASS periodically reports rental rates negotiated on an areal basis in addition to the AUM basis. When these records exist together, they can be used with RAP-V2 production data to calculate UF as shown in ***equation 8*** (a full derivation is provided in the supplemental text). This approach assumes that the value of herbaceous production is comparable in the two rental schemes.

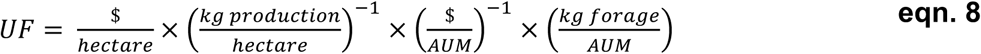

Using this method, we calculated a mean UF of 0.4011 (standard deviation = 0.0619, n = 168), slightly more conservative than the literature-based value. Importantly, military and protected non-grazing lands (e.g., National Parks) were excluded from the monetization calculations, but other federal grazing lands (i.e., BLM and USFS allotments) were included. Finally, dollar estimates were inflation-corrected to 2019 dollars using the consumer price index from the U.S. Bureau of Labor Statistics.

### 2.5 Evaluation of model uncertainty and error

To assess uncertainty in our yield gap estimates, we used Mean Absolute Percentage Error (MAPE) to provide upper and lower bound yield gap estimates for our annual data (**equations 9 and 10**).

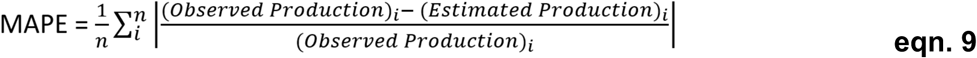

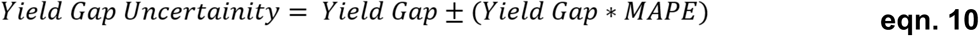

To incorporate uncertainty in the monetary estimate, we used a Monte Carlo simulation (n = 10,000) to account for variability in biomass estimates, forage rental rates ($/AUM), and UF. We assumed forage rental rates ($/AUM) varied by ± 20% (random uniform distribution) and used random normal UF estimates derived from ***equation 8***.

Uncertainty in production estimates attributable to satellite signal quality and atmospheric noise (e.g., clouds) was assessed by calculating the yield gap with and without data imputation and filtering measures. Further details of the approach and results are presented in supplemental text.

## 3. Results

Rangelands gained 77,323 ± 1,222 km^2^ of tree cover between 1990 and 2019 (**Fig. 2**). Absolute tree cover increased from 154,502 km^2^ to 231,825 km^2^ (**Table 2**). Tree cover grew on average 2,577 km^2^ per year, with annual increases in all years. Tree cover in grasslands grew by 85% over this period, with shrublands and open woodlands increasing by 43% and 40%, respectively. In total, we observed increased tree cover on 25% (684,852 km^2^) of rangeland area (pixels) between 1990 and 2019. In contrast, tree cover declined across roughly 6% of rangelands (180,461 km^2^).

**Table 2.** Absolute tree cover change from 1990 - 2019

|  | Intact Biome Area(†) | Tree Cover 1990 |  | Tree Cover 2019 |  | Tree Cover Expansion |  |
| --- | --- | --- | --- | --- | --- | --- | --- |
|  | km<sup>2</sup> | km<sup>2</sup> | % | km<sup>2</sup> | % | km<sup>2</sup> | % |
| Grasslands | 1,032,089 | 30,012 | 2.91 | 55,537 | 5.38 | 25,524 | 2.47 |
| Shrublands | 1,265,754 | 42,441 | 3.35 | 60,852 | 4.81 | 18,411 | 1.45 |
| Open Woodlands | 438,216 | 81,379 | 18.57 | 114,625 | 26.16 | 33,246 | 7.59 |
| Sparsely vegetated (‡) | 47,895 | 669 | 1.40 | 812 | 1.69 | 142 | 0.30 |
| Total | 2,783,955 | 154,502 | 5.55 | 231,825 | 8.28 | 77,323 ± 1,222 | 2.73 ± 0.04 |

Notes: † Intact biome areas calculated from Landfire BPS v1.4, NLCD 2016 and NASS data. Areas exclude row-crop agricultural lands and the built environment. ‡ Sparsely vegetated class includes all sparsely vegetated systems identified in BPS v1.4.0; these areas typically are found as subcomponents of the three primary biomes included in our analysis. The total tree data includes the 95% confidence interval based upon incorporating model error rate and trend analysis of the time-series data.

The magnitude and direction of tree cover change varied by region. Tree cover was found to be increasing in 11 of 14 regions examined, while substantial losses in tree cover were observed in parts of the desert southwest and along the Pacific Coast (**Fig. 3)**. Tree cover trends appeared to be accelerating in the central and northern rangelands, while tree cover growth appeared to slow across portions of southeast rangelands. **Supplemental Fig. 2** shows a map highlighting areas (pixels) of increasing and decreasing tree cover.

**Figure 3.**
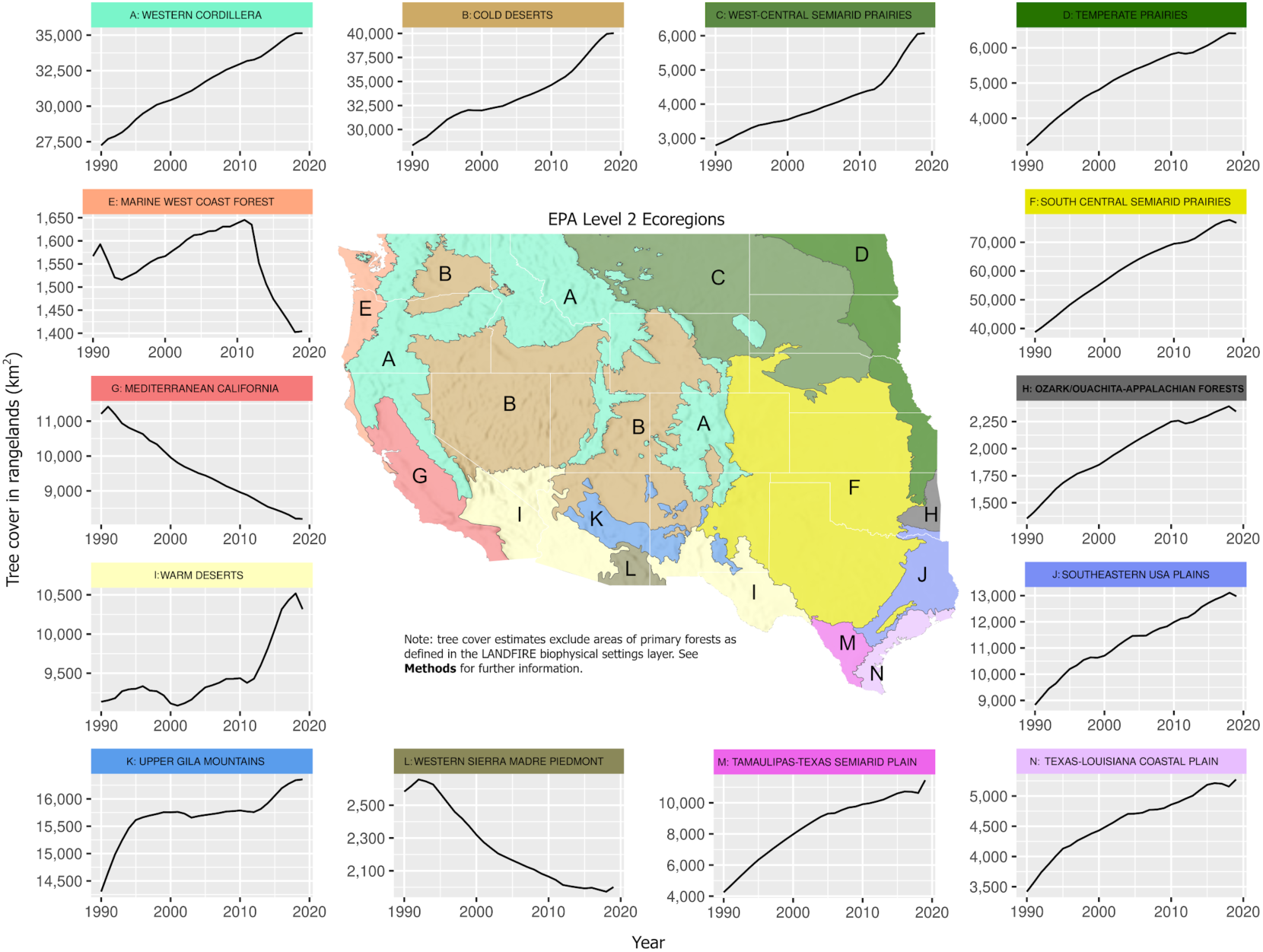
Rangeland tree cover by EPA Ecoregion Level 2 (1990 – 2019). Woody expansion is accelerating in parts of the northern Great Plains (*Region C*) and Great Basin (*Region B)* and slowing in parts of the southern Great Plains and Texas (*Regions F, J, M, and N)*. Sustained tree cover losses were found in the desert southwest (Region L) and California (*Region G)*.

Importantly, tree encroachment resulted in the loss of 147,700 km^2^ (range: 135,283 - 150,827 km^2^) of tree-free rangelands, transitioning roughly 8% of these lands into woodlands (**Fig. 2, Panel C**). Even where tree-free rangelands persist, they were increasingly vulnerable to encroachment due to their proximity to nearby tree seed sources. For example, the area of intact tree-free rangelands (rangelands greater than 200 meters from a tree seed source) declined by 15% (204,651 km^2^) over the 30 years.

Tree encroachment contributed to sizable losses in herbaceous production (**Fig. 4**). Production losses totaled 302 ± 30 Tg (dry biomass) from 1990 through 2019. In 2019, the yield gap totaled 20.0 ± 4.2 Tg, representing 5% - 6% of the potential forage production for all western U.S. rangelands (**Fig. 4b**).

**Figure 4.**
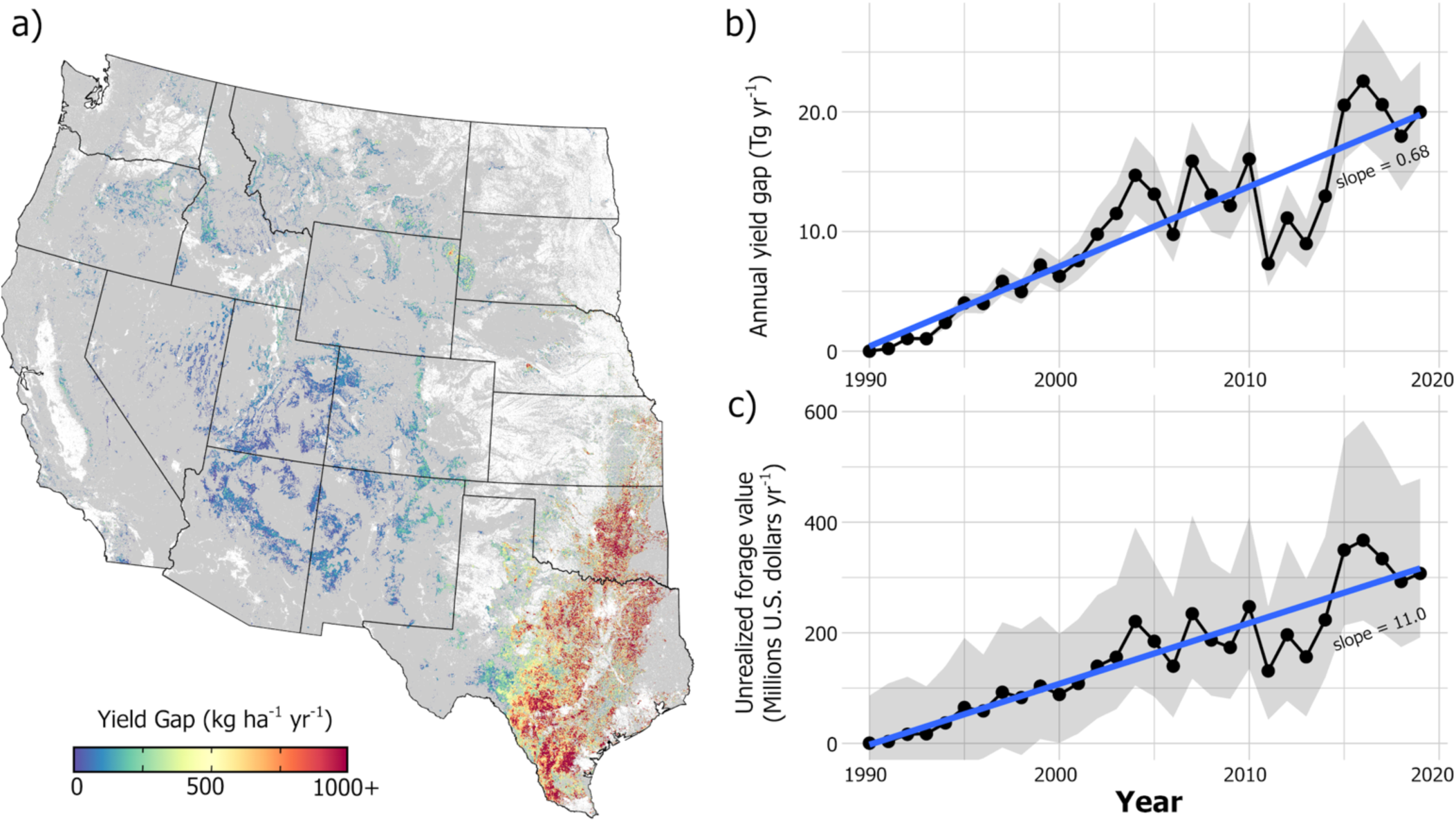
Herbaceous production yield gap attributable to woody encroachment: 1990 - 2019. a) Map of 2019 yield gap; converted agricultural lands and built environment shown in white. b) Total annual yield gap in Tg ± MAE; includes dry herbaceous biomass (grass and forb); c) The monetary value of forage lost to woody encroachment; error bars represent the 95% prediction interval.

Locally, we observed that increasing primary productivity buffers production losses during early tree colonization of rangelands. We found that herbaceous production could remain stable or even increase following the establishment and expansion of woody encroachment (**Fig. 5**). In the absence of disturbance, however, yield gaps rapidly develop as tree cover continues to increase, resulting in large net declines in production over short periods. For example, **Fig. 6** illustrates how a 30% yield gap developed in only five years following 20 years of sustained tree cover expansion.

**Figure 5.**
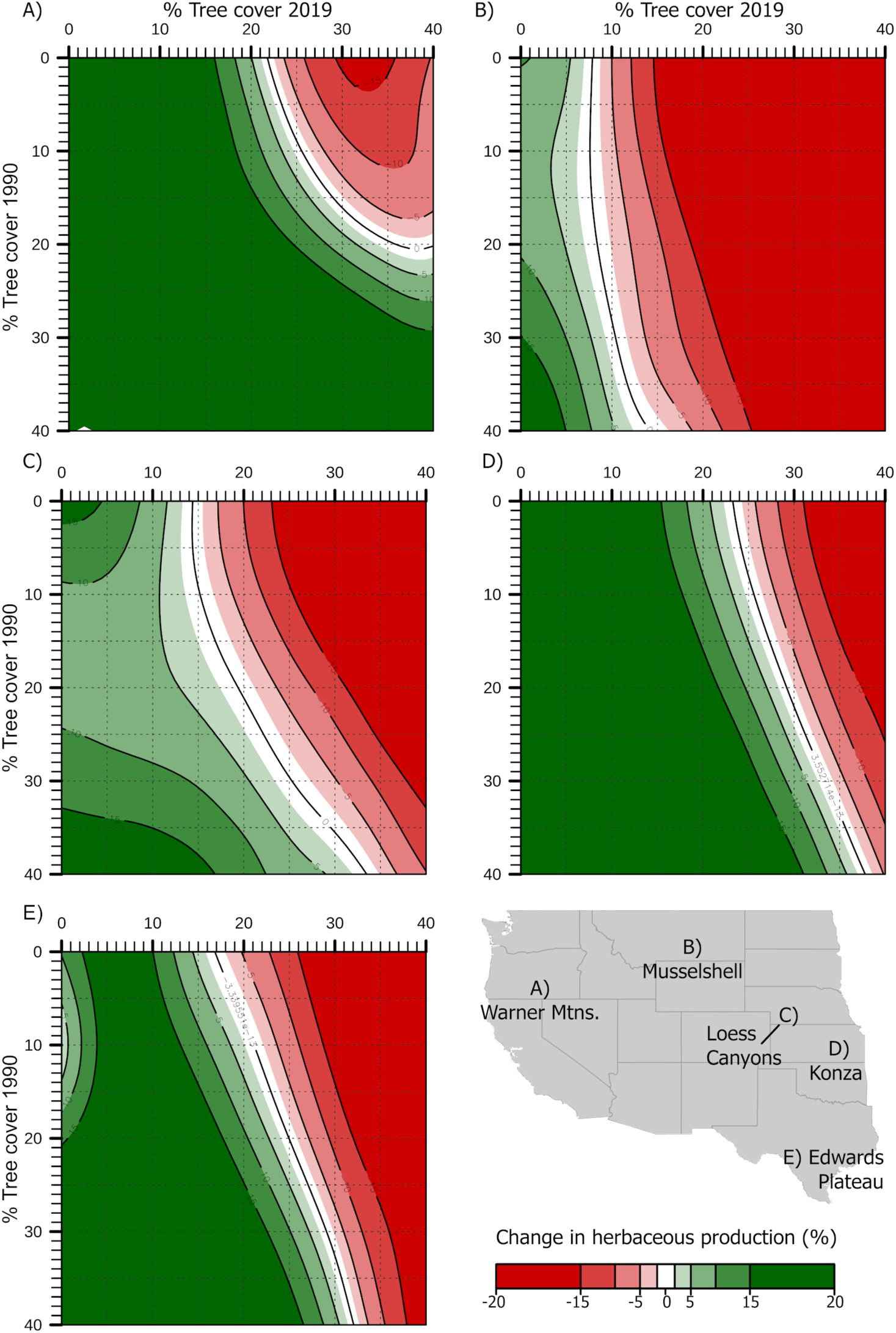
Localized relationships between tree cover change and herbaceous production from years 1990 - 2019 for five diverse locations across U.S rangelands. Impacts on herbaceous production from tree encroachment vary by location due to local biophysical conditions and climate. These data are calculated at the county scale and are truncated at ±20% to emphasize dominant threshold behavior.

**Figure 6.**
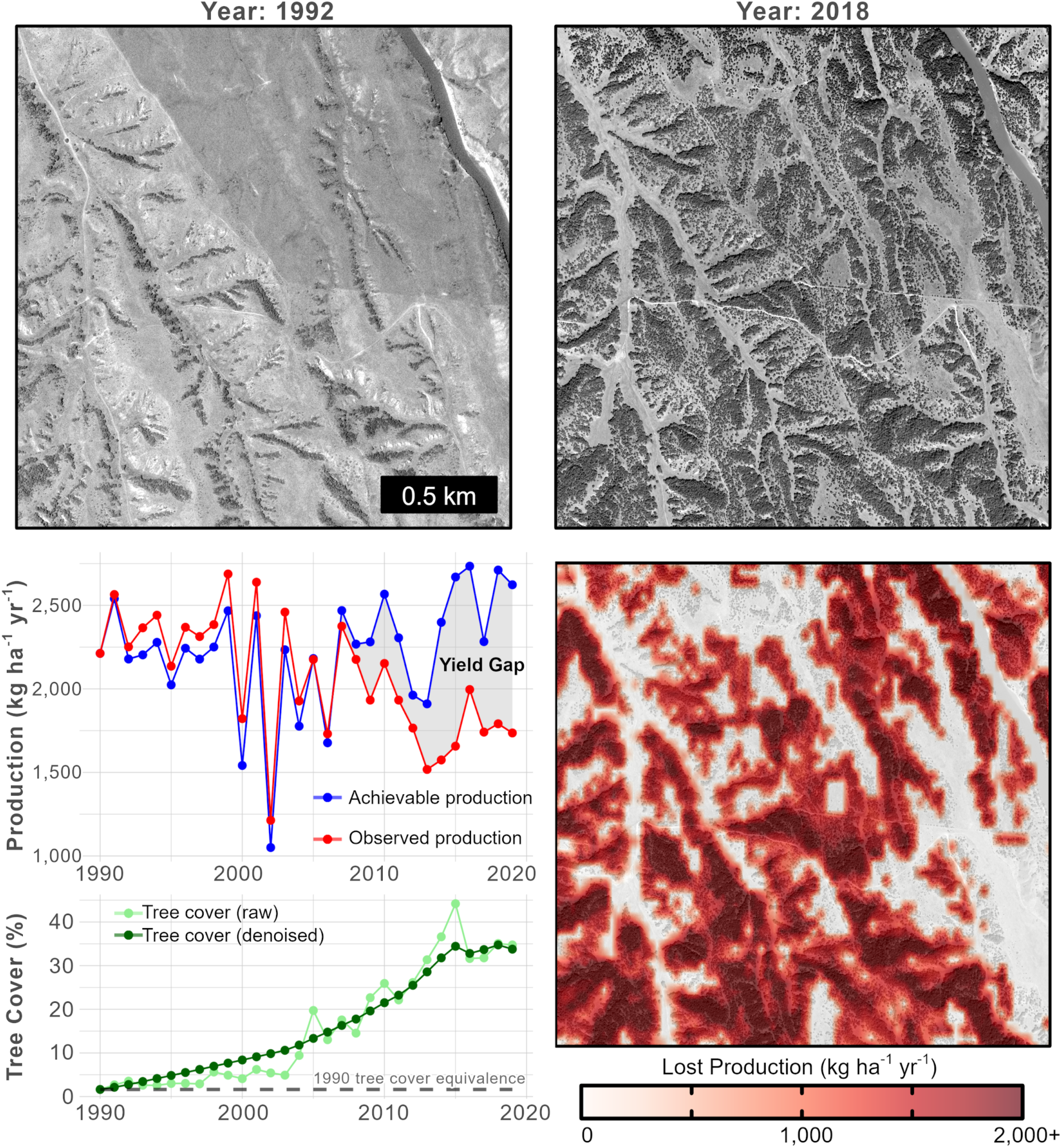
Local-scale evaluation of tree encroachment and yield gap development in the Loess Canyons, Nebraska USA (W100.44° N40.95°). Herbaceous production decreased by 34% percent between 1990 and 2019. **(Top)** Time series aerial imagery shows the dramatic expansion of Eastern Red Cedar (*Juniperus virginiana)*. (**Bottom**) Time-series and spatial trends in achievable and observed forage production show the forage yield gap’s evolution as a function of tree encroachment.

On an inflation-corrected basis, the value of herbaceous production lost to woody encroachment totaled U.S. $4.84 ± 0.72 billion over the 30 years analyzed; losses for 2019 were estimated to be $307.2 million ($192.6 - $480.3 million). Our monetary estimates were sensitive to biomass modeling error and under constrained economic variables (i.e., uncertainty in rental rates and biomass utilization). Assumptions regarding forage utilization by livestock may be particularly important but are also the hardest to constrain in space and time.

Our yield gap results were generally robust to errors introduced from satellite signal degradation and processing, particularly for years after 2000 (**Supplemental Fig. 3**). Variability in yield gap calculations attributable to signal and processing errors was approximately 3% in 2019 (yield gap range: 19.7 to 20.3 Tg). Prior to 2000, satellite data processing relied heavily on gap-filling methods. However, the imputation of these data had little impact on the interpretation of temporal trends.

## 4. Discussion

Our analysis shows that the conversion of grasslands to woodlands is widespread among rangelands of the U.S., with estimated impacts to an area roughly the size of the country of Germany (calculated as the sum of lost and vulnerable tree-free rangelands, 352,261 km^2^). These findings are consistent with model-based projections of climate-driven woodland expansion and impacts from a century of wildland fire suppression (Bond, 2008; Klemm et al., 2020; Ratajczak et al., 2014). Modest increases in tree density are well-known to have outsized impacts on ecosystem services and biodiversity in rangelands. For example, many grassland and shrubland obligate bird species are lost from rangelands when tree cover exceeds more than a few percent (Baruch-Mordo et al., 2013; Herse et al., 2018; Lautenbach et al., 2017). These results further reinforce scientific conclusions that woody encroachment is a primary change agent across broad regions of U.S. rangelands (Engle et al. 2008; Wilcox et al. 2022).

The pace of tree encroachment is similar in magnitude to recent row-crop conversion, another primary threat to the conservation and sustainability of U.S. rangelands. From 2008 to 2016, conversion of intact rangelands to cropland accelerated rapidly across the western U.S; annual conversion rates were observed to be 1,649 to 4,385 km^2^ annually (median = 2,777 km^2^, **Supplemental Fig. 4**) (Lark et al., 2020). In comparison, the median annual loss of rangelands to tree encroachment was 1,899 km^2^ over this same period. Together, these data suggest that rangelands of the western U.S. are being lost at a rate of 1,282 hectares (3,168 acres) per day, losses that are 68% higher than estimates based solely on row-crop conversion. Importantly, not all areas are experiencing tree encroachment. Large tree cover declines were observed in the southwest of our study area, attributable primarily to long-term drought (**Fig. 3, Supplemental Fig. 4)**.

We found that tree encroachment creates sizable yield gaps in herbaceous production across western U.S. rangelands. This production mediates critical ecosystem services such as below-ground C storage and is the cornerstone of food and fiber production in rangelands (Jackson et al., 2002). In an agricultural context, the annual 5% - 6% yield gap observed in this analysis is comparable to sector-wide yield losses sustained to commodity crops under extreme drought events (Lesk et al., 2016). Given that 25% of U.S rangelands are seeing increasing tree cover, the continued expansion of yield gaps this century could dramatically compound yield losses under intensifying drought cycles resulting from climate change.

Rapidly accelerating woody encroachment and production losses in the northern Great Plains are particularly concerning to working lands conservation (**Fig. 3**). This area is central to ongoing efforts to protect the North American grassland biome. Here, production losses resulting from tree encroachment may further exacerbate economic harm to small ranching operations, promoting accelerated conversion of grasslands to alternative land use types (Anadon et al., 2014; Haggerty et al., 2018; Lark, 2020). Losses of these intact grasslands threaten the last remaining large-scale migrations of native ungulates (*Antilocapra americana*) in the contiguous U.S. (Joly et al., 2019). Further, the area is critical habitat for rapidly declining obligate grassland bird populations which are particularly sensitive to habitat fragmentation and tree encroachment (Brennan & Kublesky, 2005; Rosenberg et al., 2019).

Importantly, primary production is not fixed or constant in an era of global environmental change. We observed sizable increases in herbaceous production among rangelands where tree cover remained constant (averaging 195.7 kg ha^-1^ yr^-1^ more production in 2019 than in 1990), consistent with other studies showing increased productivity across large swaths of the North American continent (Boone et al., 2018; Hicke et al., 2002; Reeves et al., 2020). These increases are attributable to increases in precipitation and growing season length as a result of climate change. Consequently, many areas undergoing early stages of woody encroachment may experience little to no loss in total herbaceous production (**Fig. 5, Supplemental Fig. 4**).

These findings illustrate both challenges and opportunities to address tree encroachment in U.S. rangelands. The lag between when trees colonize and when yield gaps are detectable may incentivize some land managers to accommodate tree encroachment until it reaches more advanced stages. In these cases, managing encroachment becomes more expensive and complex due to removing additional tree biomass and established seed stores (Roberts et al., 2018). Conversely, the time lag provides an opportunity to implement cost-effective management to prevent further tree encroachment, productivity decline, and economic losses to agricultural producers.

Removing trees in rangelands presents trade-offs between competing ecosystem services and biodiversity goals, and the scale and severity of these trade-offs should be considered by applied ecologists. In some areas, such as equatorial grasslands, tree encroachment can enhance local provisioning of shade, fuelwood, and food and also mitigate climate change impacts to communities (Linders et al., 2021). Similarly, removing encroaching trees may unintentionally impact wildlife that utilize newly developed woodland patches (Tack et al., 2022). In the context of U.S. rangelands, preventing further tree encroachment is a key biome-level strategy to protect remaining intact grassland and the obligate species that depend on tree-free landscapes (Scholtz & Twidwell, 2022).

Yield gap estimates here are consistent with other studies that investigated tree-grass interactions, including field measures from the Konza Long Term Ecological Research Site (**Supplemental Fig. 5)**. While there is consensus that herbaceous production declines under moderate to high tree cover, there is less agreement on how incipient tree encroachment impacts production (Anadon et al., 2014; Fuhlendorf et al., 2008; Scholes, 2003). We did not find evidence for declining herbaceous production during early tree colonization. However, our model-based findings should not be viewed as definitive due to interactions between shifting baseline production and model sensitivity to sparse tree cover (see **Supplemental Text**).

Our monetary estimates for lost revenues should be considered preliminary as they do not incorporate economic, policy, and social considerations that drive market valuation and the adoption of management alternatives. Locally, our estimates for forage value agree well with independent agricultural economic data (**Supplemental Table 3)**. Still, more sophisticated analyses will be required to assess how managing tree encroachment impacts ranching economics.

The loss of rangeland production and corresponding impacts to ecosystem services are often overlooked in the face of advocacy efforts calling for the afforestation of rangelands (Bastin et al., 2019).Increasing tree cover in rangelands may not benefit global climate change mitigation strategies due to biogeophysical feedbacks (Friedlingstein et al., 2019; Nuñez et al., 2021). Tree encroachment can increase C storage in rangelands (Asner et al., 2003; Connell et al., 2020), but these C storage gains are typically offset by reducing land-surface albedo at mid- to high-latitudes (Bala et al., 2007; Betts et al., 2007). For example, across the northern Great Plains of North America, tree cover would need to reach 95+% to contribute to climate cooling; such a scenario would result in the wholescale collapse of the northern Great Plains and its biodiversity therein (Hoekstra et al., 2004; Mykleby et al., 2017). The most effective climate mitigation strategy for rangelands remains preventing grassland conversion to row-crop agriculture and ensuring the massive C reservoirs found in rangeland soils are not lost to the atmosphere by land-use conversion (Bossio et al., 2020; Hoekstra et al., 2004).

## 5. Conclusion

Identifying yield gaps and mitigating the impacts of forage losses provides a new approach to support the economic sustainability of U.S. rangelands and slow land-use conversion. Herbaceous production is the cornerstone of food and fiber economics in working rangelands and evaluating it like other agricultural commodities provides a means to stack market incentives with other conservation subsidies. Implementing landscape-scale conservation strategies to close and prevent yield gaps will help maintain the function and biodiversity of rangeland ecosystems, ultimately benefiting nature and society.

## Acknowledgments

This work was made possible by the USDA-Natural Resources Conservation Service’s (NRCS) Conservation Effects Assessment Project - Wildlife Component, the NRCS’s Working Lands for Wildlife, the Arkansas Game & Fish Commission through cooperative agreement 1434-04HQRU1567, and the Bureau of Land Management. NVIDIA provided hardware used in this analysis through their Academic Hardware Grants program. The findings and conclusions in the publication are those of the authors and should not be construed to represent any official USDA or U.S. Government determination or policy. Any use of trade, firm, or product names is for descriptive purposes only and does not imply endorsement by the U.S. Government. The authors declare no conflicts of interest.

