## Supplemental Materials for "Herbaceous production lost to tree encroachment in United States rangelands"

**This PDF file includes:**

Supplementary text
Figures S1 to S6
Tables S1 to S4
SI References

**Model evaluation**

Direct comparison of the yield gap modeling results to field (ground-truth) data is challenging due to the mismatch in spatiotemporal scales between our simulation results and field measurements, plus the relative scarcity of field data that jointly track long-term tree cover expansion and herbaceous production. Previous publications (Allred et al., 2021; Jones et al., 2021) evaluated the underlying herbaceous production and tree cover models used in this analysis; the *Rangeland Analysis Platform* (RAP) website (*https://rangelands.app/products/*) provides additional information regarding the performance of these components. Here, we compare the modeling results and calculations from this analysis to herbaceous production data compiled by the USDA-NRCS (Soil Survey Staff, 2020), longitudinal production data from the Konza LTER (Blair & Nippert, 2020; Connell et al., 2020), and economic data on herbaceous production compiled by Kansas State University (Kansas State University, 2019).

Overall, we emphasize that our findings are congruent with literature-based evaluations of tree expansion impacts on herbaceous production. These studies overwhelmingly indicate that herbaceous production declines in response to moderate and high tree cover increases (Archer et al., 2011; Fuhlendorf et al., 2008; Jameson, 1967; Mitchell & Bartling, 1991; Richard Teague et al., 2008; Scholes, 2003; Scholes & Archer, 1997). The primary difference between our modeling results and the literature consensus is the relationship between tree cover and herbaceous production during the early phases of tree canopy expansion.

Our analysis showed that the herbaceous production response to tree cover expansion generally followed time-lag or threshold behavior and not inverse linear or exponential trends commonly reported in the literature (see **Fig. 5 and 6** in *main text* and discussion in Scholes, 2003 and Scholes and Archer, 1997). These differences could be attributable to one or more of the following factors. First, increasing primary productivity at the continental scale can obfuscate the relationship between tree cover and herbaceous production through time, particularly at low to moderate tree covers. Second, our analysis tracks changes in tree cover and production through time whereas most published results from U.S. rangelands rely on either 1) space-for-time substitution or 2) inverse analysis of herbaceous production following tree removal. These approaches cannot account for hysteresis effects, so they would likely not mirror our observations. Third, our model may not have enough sensitivity to disambiguate the role of tree cover and regional-scale primary productivity trends where tree cover is less than 10%. For example, we regularly see that *observed production* marginally exceeds *achievable production* as tree cover increases from 0 - 10% (e.g., **Figure 6** in *main text*). While these observations are universally within the model margin-of-error (+/- 1 MAE), they suggest that our model framework can under-estimate achievable production to a small extent when tree cover is low. However, even when our model underestimates achievable production, it does not appreciably change the interpretation of our analysis: higher achievable production during early tree invasion would only serve to increase yield gap estimates in our analysis.

**Comparison with field data from Konza LTER**: We compared our yield gap estimates to long-term production data collected from watersheds 001D and 020B at the Konza LTER (Blair & Nippert, 2020). The paired Konza watersheds are used for long-term biogeochemical experiments to study the fire and grazing impacts on nutrient and carbon cycling[56]. Watershed 001D was burned annually from 1978 - 2015 while watershed 020B was last burned in 1991. Woody cover in watershed 001D remained unchanged at less than 5%, while woody cover increased from roughly 5% to 50% in watershed 020B from 1990 to 2018; see **Figure 9** in Connell et al., 2020 for details. **Figure S6** contrasts herbaceous production at Konza to Yield Gap model estimates. While not a perfect comparison, it shows generally strong agreement between production changes in watershed 020B and our yield gap model. Both the field data and model data show dramatic increases in the production gap and yield gap between 2010 and 2015. Moreover, the production gap between watersheds 001D and 020B grew by roughly 1,600 kg/ha from the 1990s to the late 2010s. In comparison, the modeled yield gap for watershed 020B grew to 1,350 kg/ha over the same period.

Note that our model estimated that tree cover increased from 1% to 33% between 1990 and 2019 in watershed 020B. This is slightly lower than the change in woody cover calculated from the Konza field data. This discrepancy could result from shrub inclusion in the field-based woody cover estimate and account for the model’s lower yield gap estimates. Notably, the model does not account for shrub cover impacts on herbaceous production, so it will generally produce a conservative estimate for the yield gap where shrubs contribute to woody encroachment and herbaceous production declines.

***Comparison against SSURGO data:*** Achievable production estimates from our modeling estimates are within the *normal* and *favorable* forage production distributions predicted from the USDA NRCS SSURGO database (**Figure S6**). These comparisons indicate that the model’s achievable production estimates are similar, but relatively conservative relative to production estimates developed by the USDA NRCS, particularly under high production scenarios. These findings are consistent with the previous analysis of production values from the RAP dataset compared to SSURGO and NRI data (Jones et al., 2021). Importantly, the values from the NRCS in both the SSURGO and NRI databases should be considered estimates, rather than empirical data, as they incorporate subjective estimations and correction factors.

The degree to which lower production estimates may impact our yield gap estimates is not fully resolved, but it does not appear to contribute to an overestimate of yield gaps in our analysis. For example, our comparison with Konza field data showed our model estimates for yield gap were ~ 15% lower than what field measures would estimate, even when some of the annual production data differed by as much as 40% between the remote sensing and field-based measures of production.

**Comparison to independent economic data**: We compared our forage value estimates to independent valuation estimates produced by the Kansas State Agricultural Economics department in cooperation with the Kansas Department of Agriculture (Kansas State University, 2019). The area covered in that analysis included 14 counties for eastern Kansas which make up the majority of the area with Tall Grass Prairie. We estimated the area of grazing lands in these 14 counties to be approximately 1.98 million hectares producing 2,970 Gg of forage. Both calculations use the herbaceous production yield gap from our model, so cannot be considered completely independent; calculations using the Kansas State data are only sensitive to the relative yield gap from our model (roughly 14%) , not the absolute values of achievable and observed herbaceous production.

The value of forage in our model used state-level data and was calculated using ***equation 7*** in the main text. The forage value from the Kansas State economic analysis is calculated using **equation S1**. Herbaceous production for this area averaged 1494.6 kg per acre and the herbaceous production yield gap was calculated to be 491.6 Gg.

$$\frac{\$}{kg\ production} = \frac{\$}{Acre} * \frac{Acre}{kg\ production}$$

**eqn S1**.

We found that the estimates from the two economic calculations differed by 21% (**Table S3**), showing reasonable agreement between the state-level data used in our model and the local data collected by the Kansas State Agricultural Economists.

**Model Sensitivity to satellite measures and productivity modeling.**

Herbaceous production in our model relies on the Landsat-based estimates of net primary productivity calculated using a modified MOD17 algorithm (Robinson et al., 2018; Running et al., 2004). Atmospheric effects, cloud cover, data transmittal, and sensor errors can all contribute to signal loss and noise in these modeled data. To minimize the impact of these artifacts and temporal noise on the primary productivity product, the Landsat NPP algorithm imposes an iterative Interpolation for Data Reconstruction method to smooth the data signal (Julien & Sobrino, 2010). Additionally, gaps in the data from sustained cloud cover are filled (when possible) using climatology data from adjacent observations and years. Together, these methods provide a statistically robust way to impute missing or poor-quality data. However, these methods can also contribute to some variability in the satellite-driven productivity estimates based on the degree of data smoothing and imputation in each pixel.

To evaluate these uncertainties, we evaluated our yield gap estimates based on three data imputation levels imposed at the pixel-level in our datasets (see Robinson et al., 2018 for details on the gap-fill model).

1) Strict filtering: No imputation or filtering. Pixels with low quality or missing data are masked out of the analysis.

2) Gap-filled data: Standard gap-filling technique imposed on the Landsat NPP dataset with gap-filling from climatology data and data smoothing. This processing was used in our analysis.

3) No filtering: Use raw data without gap-filling, smoothing, and imputation.

The results of this sensitivity analysis are shown in **Fig. S3**. Overall, we found that the underlying signal quality from the Landsat sensor had a relatively small impact on our yield gap estimates for years 2000 - 2019. Satellite Observations for the first decade (1990 - 2000) relied heavily on the gap-filling algorithm, attributable primarily to the absence of suitable climatology data to correct the data and particularly cloudy conditions in the southern Great Plains during the growing season. For 2019, the yield gap estimates ranged from 19.7 Tg to 20.3 Tg across the three methods, a relative difference of only 3%.

**Derivation of *utilization factor* (equation 8 in the main text).**

To calculate the dollar value of herbaceous production from AUM rental rates requires using a utilization factor (UF). UF represents the fraction of total herbaceous production consumed as forage by livestock (**equation S2**).

$$UF = \frac{kg\ forage}{kg\ production} \qquad \textbf{eqn. S2}$$

A UF is needed because not all herbaceous production is consumed by animal livestock during grazing. Grazing leases in the US are most commonly negotiated by AUM, which is a measure of forage consumed by livestock rather than a direct measure of pasture production. Using a UF conversion provides a means to convert rental rates from the basis of consumed forage to total production.

Grazing operations in U.S. agriculture generally abide by the principle of use-half leave-half when prescribing grazing intensity. Holechek et al. (2010) estimated UF as 0.43 under moderate grazing intensity, but other spatially or temporally explicit data is generally sparse.

The calculation used in our analysis to estimate the monetary value of herbaceous production is presented in **equation S3**.

$$\frac{\$}{kg\ production} = \frac{\$}{AUM} \times \left(\frac{kg\ forage}{AUM}\right)^{-1} \times UF \qquad \textbf{eqn. S3}$$

Where dollars per Animal Unit Month (*$/AUM*) represents the grazing rental rate, and *kg forage / AUM* represents the mass of consumed forage per AUM.

In addition to reporting rental rate information based on AUM, the USDA also intermittently reports rental rates on an areal basis ($ / area). Using the areal rental rate to estimate the monetary value of herbaceous production is more direct than the calculation presented in **equation S3.** However, areal rental rates are not reported for every state and year in our analysis, so we cannot use them directly in our calculations. **Equation S4** shows the estimate of production value based on areal rental rates ($/hectare) from USDA and areal production estimates from the Rangeland Analysis Platform.

$$\frac{\$}{kg\ production} = \frac{\$}{hectare} \times \left(\frac{kg\ Production}{hectare}\right)^{-1} \qquad \textbf{eqn. S4}$$

In cases where both areal and AUM rental rates are available for a given state and year, we can substitute **equation S4** into **equation S3**, as shown in **equation S5**. We can then solve for UF in **equation S6**.

$$\frac{\$}{hectare} \times \left(\frac{kg\ Production}{hectare}\right)^{-1} = \frac{\$}{AUM} \times \left(\frac{kg\ forage}{AUM}\right)^{-1} \times UF \qquad \textbf{eqn. S5}$$

$$UF = \frac{\$}{hectare} \times \left(\frac{kg\ Production}{hectare}\right)^{-1} \times \left(\frac{\$}{AUM}\right)^{-1} \times \left(\frac{kg\ forage}{AUM}\right) \qquad \textbf{eqn. S6}$$

This approach assumes that biomass value is roughly comparable in the two rental schemes. The main benefit of this approach is that it allows us to calculate a mean and variance for UF, so we’re not dependent on a single literature-based utilization factor.

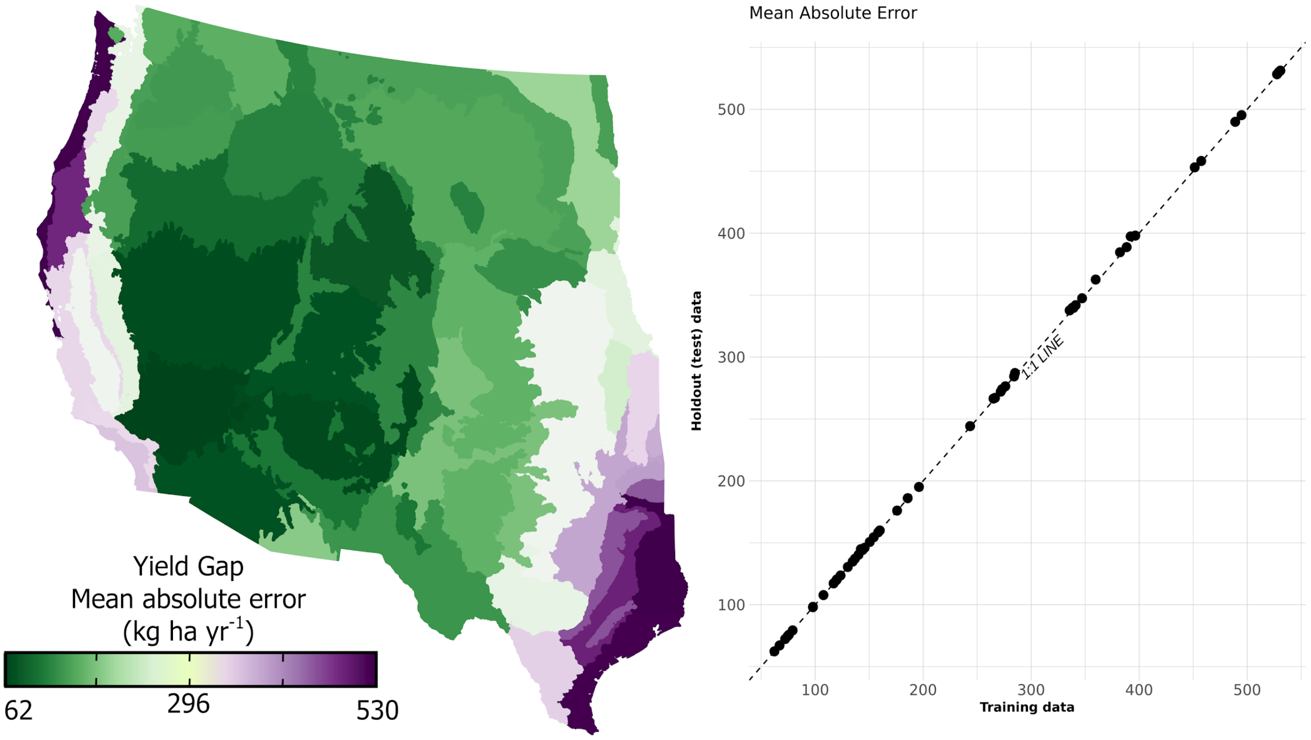


**Figure S1.** Yield gap modeling error. Left panel: average mean absolute error (MAE) for each of the Ecoregion Level 3 Models trained in this analysis. Yield gap is only tallied in our analysis when the estimated yield gap exceeds the MAE for three consecutive years; see Methods for details. Right panel: MAE values for training vs hold-out data show close agreement showing the model was not overfit.

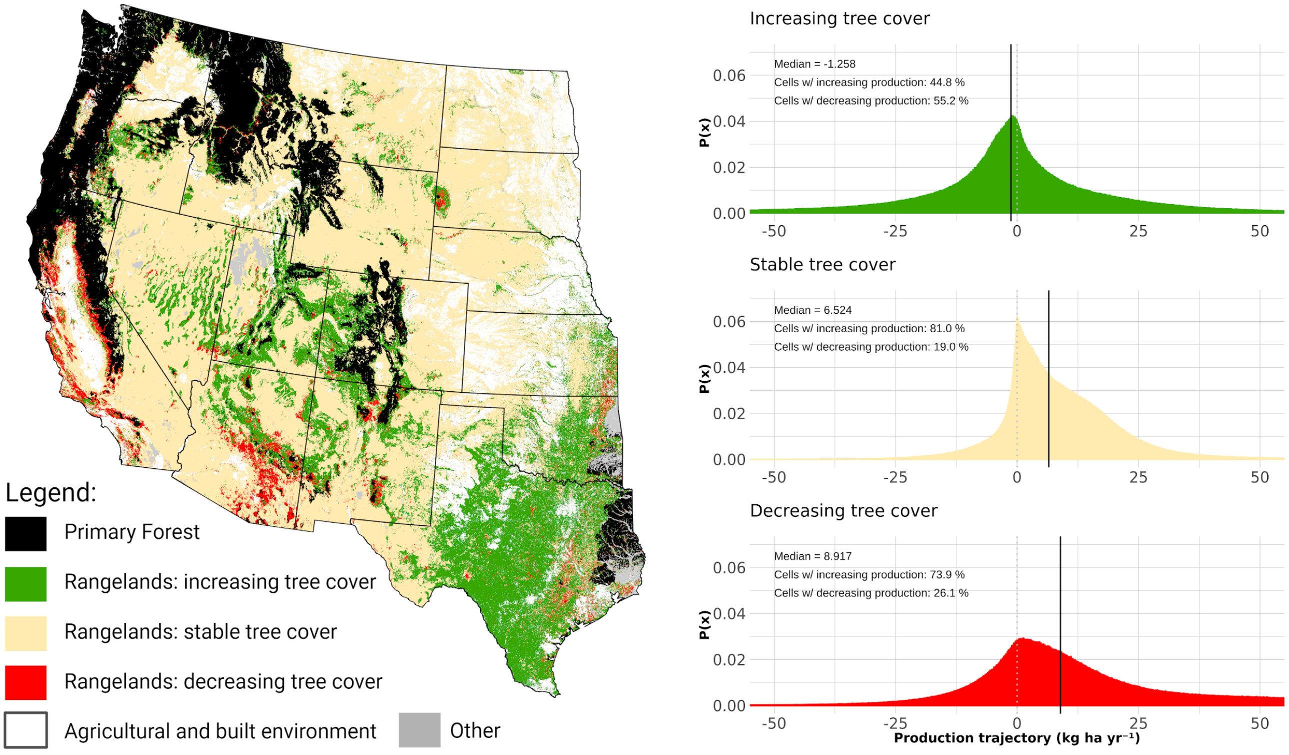


**Figure S2.** Rangeland production trends for areas with increasing, stable, and decreasing tree cover. Increasing productivity across the western United States over the past 30 years offsets production losses attributable to tree cover expansion. In many cases, absolute declines in production attributable to woody expansion occur only after substantial increases in tree cover (see Main Text and Figure 5).

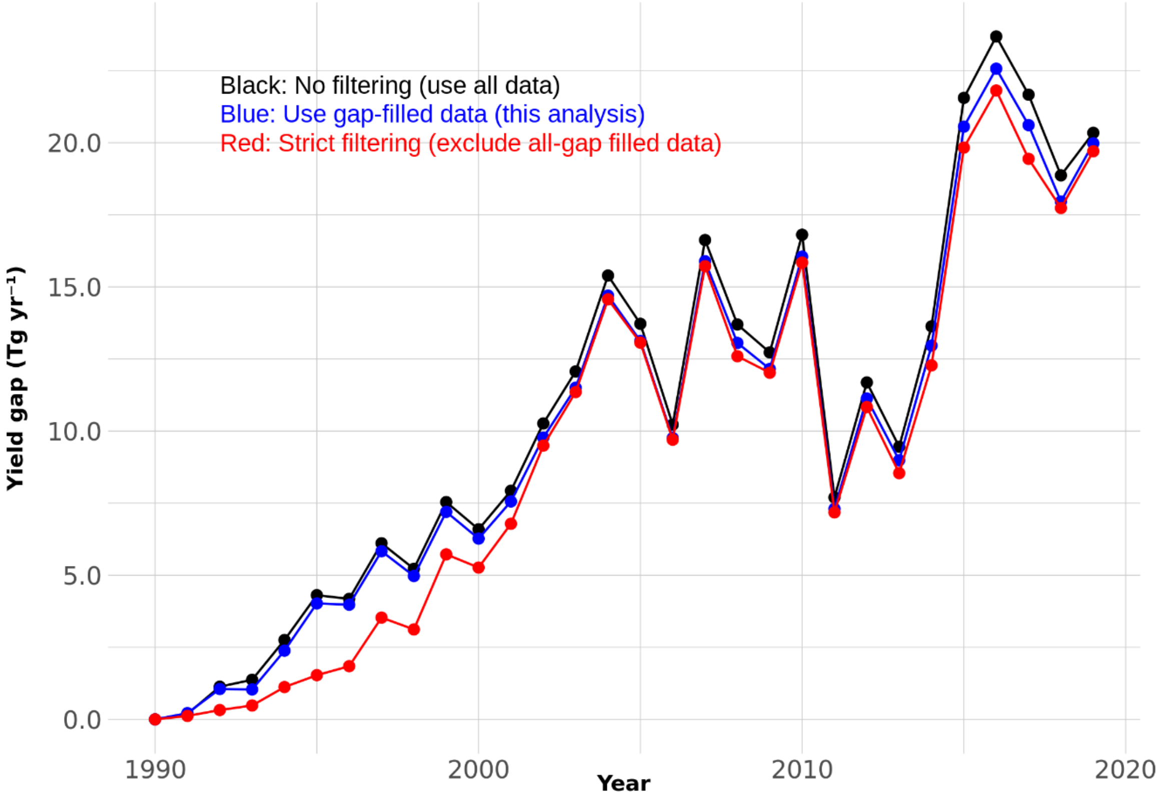


**Figure S3.** Sensitivity of yield gap estimates to uncertainty in the underlying productivity model resulting from satellite signal and processing errors.

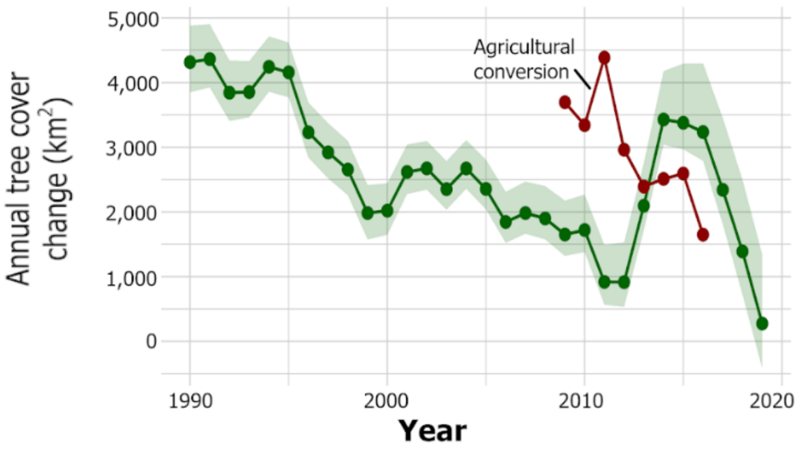


**Figure S4.** Annual net tree cover change; error bands represent the 95% prediction interval. The red time-series represents the rate of agricultural conversion for 2008 - 2016 from reference 23. Prior to 2010, agricultural conversion rates were substantially lower across the United States[67], but spatial data for the western U.S. was not available. The lower rate of tree cover expansion observed from 2016 through 2019 is primarily attributable to tree cover declines along the west coast (**see Figure S4**).

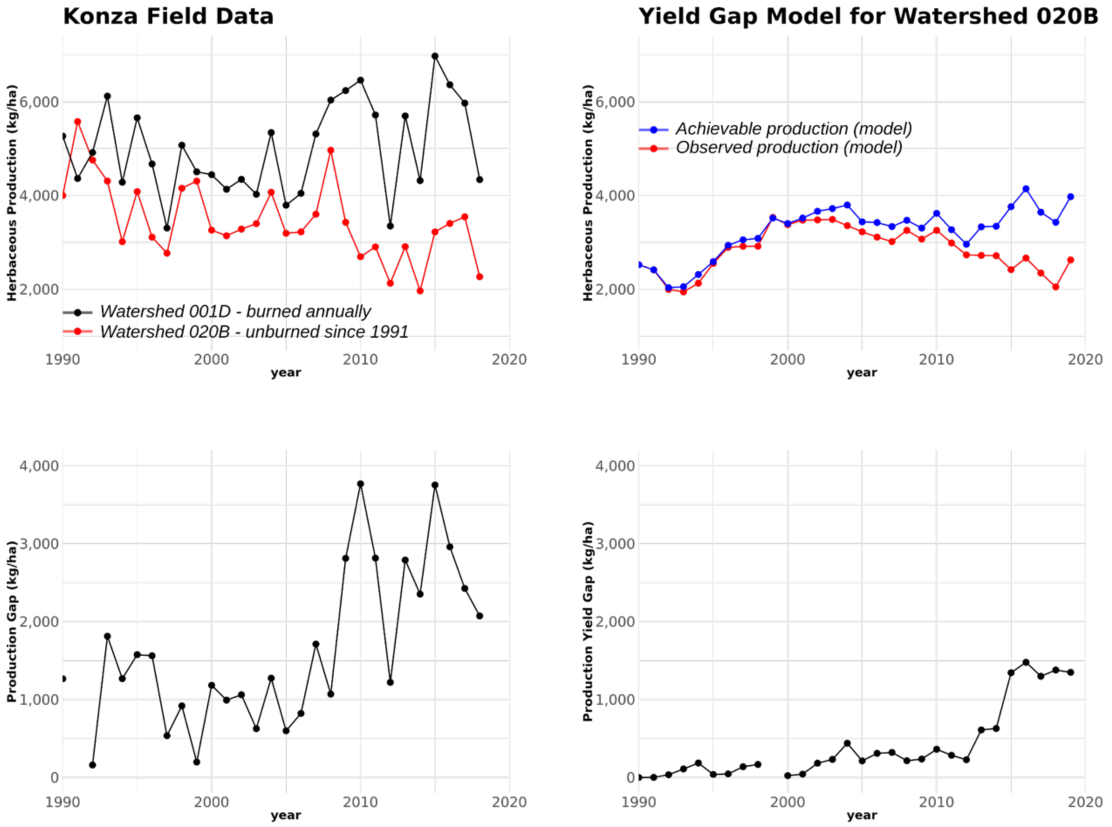


**Figure S5.** Comparison between field data from Long-Term Ecological Research (LTER) Network - Konza (left panels) and yield gap model (right panels). The right panels show production and yield gap estimates for wasterhsed 020B. Actual production for Watershed 020B is shown in red in the top panels. The production gap between the burned and unburned plots at Konza grew by roughly 1,600 kg/ha as tree cover increased. In contrast, the yield gap model estimated a yield loss of 1,300 kg/ha for the same location.

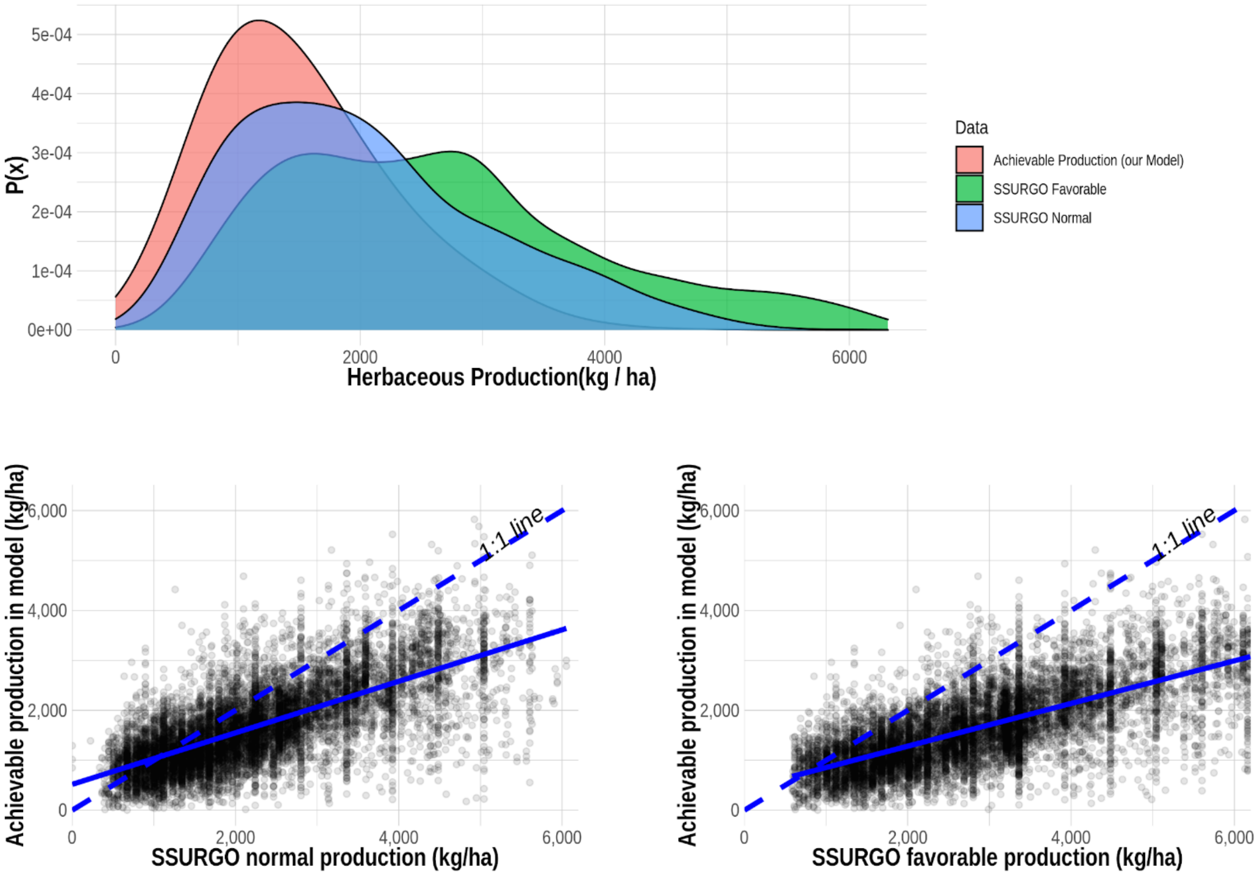


**Figure S6.** Achievable production values estimated in the model provide reasonable and conservative estimates of production compared to USDA production data. SSURGO normal represents USDA estimates for annual average production. SSURGO favorable represents production estimates under a highly productive year. These results show that our yield gap modeling framework does not produce unreasonably high estimates of production when tree cover is modified.

| **Table S1**. LandTrendr Parameters | |
|---|---|
| Parameters | Value |
| Maximum segments | 5 |
| Spike threshold | 0.9 |
| Vertex count overshoot | 3 |
| Prevent one-year recovery | TRUE |
| Recovery threshold | 0.25 |
| p-value threshold | 0.05 |
| Best model proportion | 0.75 |
| Minimum observations needed | 30 |

**Table S2**. XGBoost Parameters

| Parameter | Value |
|---|---|
| nfolds | int 0 |
| keep_cross_validation_models | logi TRUE |
| keep_cross_validation_predictions | logi FALSE |
| keep_cross_validation_fold_assignment | logi FALSE |
| score_each_iteration | logi FALSE |
| ignore_const_cols | logi TRUE |
| stopping_rounds | int 0 |
| stopping_tolerance | num 0.001 |
| max_runtime_secs | num 0 |
| seed | int 1234 |
| distribution | chr "gaussian" |
| tweedie_power | num 1.5 |
| categorical_encoding | chr "OneHotInternal" |
| quiet_mode | logi TRUE |
| ntrees | int 60 |
| max_depth | int 15 |
| min_rows | num 1 |
| min_child_weight | num 1 |
| learn_rate | num 0.3 |
| eta | num 0.3 |
| sample_rate | num 1 |
| subsample | num 1 |
| col_sample_rate | num 0.8 |
| colsample_bylevel | num 0.8 |
| col_sample_rate_per_tree | num 1 |
| colsample_bytree | num 1 |
| colsample_bynode | num 1 |
| max_abs_leafnode_pred | num 0 |
| max_delta_step | num 0 |
| score_tree_interval | int 0 |
| min_split_improvement | num 0 |
| gamma | num 0 |
| nthread | int -1 |
| build_tree_one_node | logi FALSE |
| calibrate_model | logi FALSE |
| max_bins | int 256 |
| max_leaves | int 0 |
| sample_type | chr "uniform" |
| normalize_type | chr "tree" |
| rate_drop | num 0.1 |
| one_drop | logi FALSE |
| skip_drop | num 0.5 |
| tree_method | chr "hist" |
| grow_policy | chr "lossguide" |
| booster | chr "dart" |
| reg_lambda | num 1 |
| reg_alpha | num 0 |
| dmatrix_type | chr "sparse" |
| backend | chr "cpu" |
| gpu_id | int 0 |
| gainslift_bins | int -1 |

**Table S3.** Comparison of economic estimates from model to independent rental rates in Kansas (2019).

| | | | Forage Value | |
|---|---|---|---|---|
| **Economic Data** | **Rental Rate** | **Units** | **Produced** | **Lost (Yield Gap)** |
| This Model (USDA NASS) | 18.5 | $/AUM | $157,569,444 | $10,553,453 |
| Kansas State Bluestem Report | 25.85 | $/Acre | $126,958,073 | $8,503,210 |
| **Difference** | | | | **21.50%** |

Notes: The area included in this analysis comprises tallgrass prairie rangelands in 14 eastern Kansas counties. The total area included in this analysis was 1,987,550 hectares producing 2,970 Gg of forage. The calculated yield gap estimate for this area was 491.6 Gg - roughly 14% of achievable production. Our model assumes a utilization factor of 0.4 while the monetary estimate from the Kansas State calculation does not require a UF.

**Table S4 |** Annual state level yield gap and lost revenue data. Yield gap values are in Mg and lost revenue in thousands of dollars.

| State | Variable | 1990 | 1991 | 1992 | 1993 | 1994 | 1995 | 1996 | 1997 | 1998 | 1999 | 2000 | 2001 | 2002 | 2003 | 2004 | 2005 | 2006 | 2007 | 2008 | 2009 | 2010 | 2011 | 2012 | 2013 | 2014 | 2015 | 2016 | 2017 | 2018 | 2019 |
|---|---|---|---|---|---|---|---|---|---|---|---|---|---|---|---|---|---|---|---|---|---|---|---|---|---|---|---|---|---|---|---|
| Arizona | Yield gap | 0 | 4,871 | 19,085 | 24,964 | 43,627 | 49,579 | 52,841 | 54,591 | 41,457 | 63,042 | 56,846 | 42,914 | 48,900 | 34,367 | 36,085 | 26,065 | 45,723 | 60,985 | 27,617 | 25,963 | 10,317 | 26,494 | 7,207 | 2,043 | 13,666 | 15,527 | 21,819 | 1,686 | 40,896 | 11,045 |
| | lost Rev. | 0 | 53 | 223 | 293 | 500 | 555 | 649 | 656 | 506 | 831 | 705 | 503 | 589 | 416 | 453 | 317 | 538 | 698 | 323 | 314 | 126 | 314 | 84 | 23 | 154 | 175 | 243 | 19 | 483 | 122 |
| California | Yield gap | 0 | 9,935 | 4,652 | -8,335 | -16,419 | -33,212 | -38,980 | -32,304 | -47,490 | -67,736 | -81,791 | -85,256 | -74,061 | -111,842 | -110,361 | -108,197 | -112,643 | -113,850 | -140,383 | -145,722 | -170,316 | -140,611 | -166,747 | -177,212 | -194,912 | -209,066 | -226,854 | -248,542 | -248,050 | -267,840 |
| | lost Rev. | 0 | 208 | 99 | -178 | -362 | -679 | -744 | -633 | -994 | -1,459 | -1,733 | -1,856 | -1,563 | -2,434 | -2,513 | -2,531 | -2,735 | -2,688 | -3,443 | -3,365 | -3,800 | -3,208 | -4,180 | -4,400 | -4,885 | -4,998 | -6,449 | -6,196 | -6,564 | -6,651 |
| Colorado | Yield gap | 0 | 4,178 | 9,522 | 17,511 | 34,653 | 60,002 | 63,947 | 75,740 | 77,656 | 100,497 | 95,649 | 97,363 | 59,185 | 101,394 | 114,652 | 123,107 | 106,109 | 144,998 | 114,507 | 124,111 | 137,053 | 102,942 | 107,026 | 130,218 | 196,648 | 252,601 | 230,264 | 262,351 | 218,297 | 236,758 |
| | lost Rev. | 0 | 85 | 204 | 349 | 708 | 1,203 | 1,378 | 1,582 | 1,668 | 2,147 | 1,944 | 1,941 | 1,230 | 2,125 | 2,431 | 2,711 | 2,264 | 3,112 | 2,288 | 2,523 | 2,797 | 2,077 | 2,351 | 2,902 | 4,189 | 5,375 | 4,981 | 6,032 | 5,029 | 5,082 |
| Idaho | Yield gap | 0 | 634 | 276 | 907 | 3,175 | 11,202 | 13,636 | 16,287 | 20,882 | 20,810 | 22,584 | 19,013 | 23,062 | 28,296 | 32,433 | 44,745 | 44,665 | 35,756 | 51,688 | 44,915 | 47,928 | 54,205 | 47,502 | 42,336 | 65,784 | 89,839 | 94,241 | 97,029 | 94,287 | 88,027 |
| | lost Rev. | 0 | 14 | 6 | 17 | 62 | 220 | 263 | 313 | 410 | 411 | 424 | 366 | 445 | 547 | 621 | 850 | 841 | 706 | 897 | 783 | 782 | 1,037 | 939 | 836 | 1,360 | 1,911 | 2,097 | 2,114 | 2,005 | 1,839 |
| Kansas | Yield gap | 0 | 165 | 46,706 | 96,740 | 141,795 | 193,642 | 189,760 | 274,047 | 327,410 | 328,599 | 311,583 | 346,478 | 571,106 | 579,496 | 689,501 | 699,767 | 570,205 | 702,153 | 748,081 | 765,307 | 910,283 | 737,615 | 706,257 | 731,661 | 971,795 | 1,233,484 | 1,317,222 | 1,193,786 | 1,251,335 | 1,362,304 |
| | lost Rev. | 0 | 4 | 1,085 | 2,244 | 3,122 | 3,958 | 4,305 | 5,318 | 7,687 | 6,729 | 6,441 | 7,254 | 12,242 | 12,613 | 14,076 | 14,349 | 11,327 | 14,064 | 14,430 | 14,286 | 18,575 | 16,537 | 15,057 | 15,839 | 21,919 | 29,332 | 30,119 | 30,339 | 31,044 | 29,243 |
| Montana | Yield gap | 0 | -1,313 | -3,155 | -4,340 | -7,095 | 8,707 | 20,609 | 20,821 | 23,267 | 38,627 | 18,939 | 24,840 | 22,262 | 35,057 | 27,010 | 56,336 | 54,071 | 61,189 | 63,407 | 62,061 | 84,317 | 81,723 | 60,622 | 110,013 | 188,007 | 227,191 | 253,269 | 259,821 | 267,540 | 266,962 |
| | lost Rev. | 0 | -30 | -79 | -102 | -168 | 202 | 460 | 473 | 534 | 908 | 460 | 620 | 554 | 859 | 674 | 1,386 | 1,289 | 1,558 | 1,581 | 1,545 | 2,111 | 2,091 | 1,606 | 2,942 | 5,418 | 6,540 | 7,513 | 7,704 | 7,743 | 7,589 |
| Nebraska | Yield gap | 0 | -9,628 | -13,691 | -13,074 | -16,561 | -17,427 | -2,278 | 1,198 | 13,604 | 13,773 | 530 | 36,791 | 47,182 | 61,119 | 66,897 | 103,616 | 76,710 | 129,535 | 103,961 | 147,262 | 178,222 | 157,561 | 114,808 | 167,011 | 250,485 | 296,429 | 328,541 | 270,132 | 337,196 | 364,364 |
| | lost Rev. | 0 | -311 | -429 | -456 | -580 | -597 | -78 | 42 | 470 | 466 | 18 | 1,269 | 1,626 | 2,128 | 2,416 | 3,541 | 2,709 | 4,263 | 3,581 | 5,050 | 6,207 | 5,673 | 4,228 | 7,124 | 11,927 | 14,469 | 16,649 | 13,011 | 18,324 | 17,207 |
| Nevada | Yield gap | 0 | 125 | 157 | 1,365 | 2,732 | 9,320 | 7,954 | 8,313 | 10,020 | 9,779 | 8,561 | 7,360 | 6,992 | 7,014 | 5,994 | 7,700 | 6,464 | 6,959 | 7,079 | 5,944 | 4,928 | 2,577 | 3,984 | 4,176 | 8,657 | 8,245 | 17,543 | 22,372 | 16,049 | 12,337 |
| | lost Rev. | 0 | 3 | 3 | 25 | 48 | 160 | 132 | 138 | 166 | 157 | 140 | 123 | 121 | 119 | 100 | 143 | 124 | 129 | 132 | 90 | 84 | 44 | 77 | 70 | 125 | 111 | 217 | 268 | 180 | 143 |
| New Mexico | Yield gap | 0 | 11,536 | 18,929 | 26,517 | 61,993 | 88,499 | 100,467 | 120,694 | 113,412 | 144,567 | 136,464 | 146,873 | 111,848 | 115,347 | 152,734 | 169,702 | 180,778 | 194,332 | 166,627 | 155,609 | 186,516 | 111,026 | 138,683 | 173,473 | 199,765 | 300,625 | 288,795 | 321,514 | 269,457 | 264,456 |
| | lost Rev. | 0 | 76 | 278 | 411 | 1,003 | 1,506 | 1,685 | 1,961 | 1,816 | 2,265 | 2,116 | 2,509 | 1,623 | 1,599 | 2,327 | 2,449 | 2,660 | 3,058 | 2,525 | 2,152 | 2,918 | 1,903 | 2,240 | 2,872 | 3,379 | 5,832 | 4,997 | 6,031 | 5,093 | 4,910 |
| North Dakota | Yield gap | 0 | -197 | -236 | -2,211 | -4,661 | -4,559 | -5,254 | -7,351 | -8,207 | -6,954 | -8,333 | -6,624 | -9,211 | -9,225 | -9,929 | -9,064 | -13,568 | -10,610 | -12,334 | -7,599 | -8,878 | -2,812 | -8,489 | 4,233 | 6,760 | 9,755 | 24,580 | 11,324 | 12,955 | 12,188 |
| | lost Rev. | 0 | -4 | -5 | -45 | -91 | -91 | -105 | -126 | -152 | -128 | -156 | -114 | -190 | -201 | -203 | -189 | -289 | -243 | -268 | -168 | -219 | -67 | -197 | 97 | 163 | 250 | 638 | 260 | 375 | 255 |
| Oklahoma | Yield gap | 0 | 76,128 | 341,767 | 206,830 | 716,464 | 971,918 | 854,107 | 1,049,736 | 1,075,319 | 1,296,254 | 1,761,550 | 1,797,277 | 2,555,765 | 2,563,566 | 2,812,955 | 2,996,961 | 2,477,524 | 3,206,150 | 3,195,588 | 2,932,760 | 3,266,175 | 2,244,515 | 2,528,331 | 2,485,497 | 3,251,267 | 4,016,385 | 4,144,194 | 4,278,007 | 3,791,232 | 4,054,652 |
| | lost Rev. | 0 | 1,199 | 4,755 | 3,015 | 8,892 | 13,243 | 11,304 | 15,522 | 17,613 | 18,465 | 21,242 | 24,084 | 31,607 | 28,931 | 35,339 | 36,417 | 30,258 | 41,284 | 39,626 | 36,497 | 42,212 | 32,560 | 36,914 | 36,714 | 48,481 | 61,327 | 64,027 | 64,716 | 55,985 | 54,104 |
| Oregon | Yield gap | 0 | 1,574 | -506 | -1,733 | -8,995 | -5,337 | -2,688 | -1,157 | 2,890 | 6,226 | 6,431 | 9,165 | 11,386 | 22,308 | 25,178 | 29,833 | 27,979 | 21,780 | 28,186 | 31,443 | 40,026 | 52,961 | 45,776 | 29,175 | 49,662 | 50,519 | 47,742 | 44,204 | 47,331 | 46,219 |
| | lost Rev. | 0 | 31 | -10 | -35 | -162 | -106 | -52 | -22 | 58 | 123 | 119 | 186 | 222 | 450 | 514 | 589 | 515 | 439 | 544 | 635 | 773 | 1,034 | 887 | 557 | 1,058 | 1,043 | 1,003 | 883 | 979 | 992 |
| South Dakota | Yield gap | 0 | -2,162 | -4,396 | -2,957 | -2,857 | 8,786 | 25,102 | 36,634 | 46,824 | 72,241 | 62,445 | 74,282 | 52,137 | 60,141 | 52,851 | 77,771 | 64,862 | 78,813 | 75,303 | 72,065 | 81,836 | 70,626 | 62,838 | 71,928 | 108,466 | 120,031 | 125,435 | 102,325 | 125,039 | 130,306 |
| | lost Rev. | 0 | -60 | -116 | -76 | -75 | 238 | 626 | 948 | 1,210 | 1,891 | 1,667 | 1,953 | 1,453 | 1,677 | 1,461 | 2,174 | 1,937 | 2,368 | 2,251 | 2,282 | 2,549 | 2,254 | 2,111 | 2,555 | 4,077 | 4,807 | 4,884 | 3,901 | 4,653 | 4,460 |
| Texas | Yield gap | 0 | 115,884 | 623,899 | 684,811 | 1,409,221 | 2,605,731 | 2,617,317 | 4,121,705 | 3,169,268 | 5,034,680 | 3,728,117 | 4,889,897 | 6,211,093 | 7,848,943 | 10,629,248 | 8,685,104 | 6,032,033 | 11,144,897 | 8,411,042 | 7,728,237 | 11,037,107 | 3,597,928 | 7,280,517 | 4,999,933 | 7,528,065 | 13,708,090 | 15,471,440 | 13,540,837 | 11,315,451 | 13,046,520 |
| | lost Rev. | 0 | 2,171 | 11,767 | 12,301 | 24,682 | 46,156 | 39,588 | 68,562 | 51,910 | 71,718 | 54,590 | 69,621 | 92,176 | 107,560 | 166,920 | 124,005 | 88,759 | 170,613 | 124,000 | 112,203 | 172,673 | 68,326 | 135,457 | 88,627 | 126,876 | 222,524 | 239,032 | 204,840 | 167,094 | 189,228 |
| Utah | Yield gap | 0 | 1,213 | 4,593 | 7,391 | 15,096 | 40,463 | 33,684 | 39,168 | 42,904 | 49,402 | 54,778 | 58,364 | 46,411 | 54,445 | 57,617 | 79,395 | 63,380 | 66,560 | 73,912 | 69,505 | 72,933 | 70,859 | 60,241 | 67,365 | 96,811 | 120,890 | 130,548 | 140,831 | 131,038 | 155,596 |
| | lost Rev. | 0 | 25 | 95 | 135 | 272 | 748 | 621 | 652 | 781 | 880 | 1,019 | 1,075 | 888 | 1,018 | 1,068 | 1,399 | 1,091 | 1,228 | 1,324 | 1,249 | 1,300 | 1,234 | 1,066 | 1,244 | 1,820 | 2,421 | 2,662 | 2,812 | 2,632 | 3,250 |
| Washington | Yield gap | 0 | 112 | -222 | -1,017 | -2,714 | -2,685 | -2,448 | -2,831 | -3,941 | -2,771 | -2,428 | -2,494 | -2,103 | -1,538 | -430 | 716 | 2,251 | 228 | -298 | 1,646 | 3,544 | 5,701 | 3,736 | 1,516 | 7,236 | 9,548 | 10,492 | 8,798 | 7,074 | 8,110 |
| | lost Rev. | 0 | 2 | -5 | -16 | -45 | -44 | -40 | -47 | -72 | -49 | -37 | -38 | -33 | -28 | -7 | 11 | 32 | 4 | -5 | 25 | 58 | 90 | 58 | 26 | 122 | 155 | 182 | 149 | 121 | 136 |
| Wyoming | Yield gap | 0 | 364 | -573 | -1,446 | 318 | 14,723 | 25,909 | 26,936 | 33,851 | 57,177 | 57,295 | 58,775 | 43,222 | 65,147 | 52,296 | 74,822 | 74,293 | 79,441 | 75,224 | 77,438 | 96,832 | 85,848 | 90,298 | 97,514 | 142,897 | 208,318 | 174,558 | 187,376 | 185,362 | 189,087 |
| | lost Rev. | 0 | 8 | -12 | -31 | 7 | 324 | 539 | 597 | 733 | 1,191 | 1,204 | 1,270 | 962 | 1,407 | 1,142 | 1,682 | 1,651 | 1,750 | 1,627 | 1,713 | 2,187 | 1,993 | 2,182 | 2,322 | 3,581 | 5,475 | 4,639 | 4,875 | 4,818 | 4,937 |
| **Total** | **Yield gap** | **0** | **213,420** | **1,046,808** | **1,031,924** | **2,369,772** | **3,999,350** | **3,953,686** | **5,802,227** | **4,939,125** | **7,158,212** | **6,229,219** | **7,515,020** | **9,725,175** | **11,454,035** | **14,634,732** | **13,058,379** | **9,700,836** | **15,809,316** | **12,989,206** | **12,090,944** | **15,978,825** | **7,259,157** | **11,082,589** | **8,940,881** | **12,891,059** | **20,458,412** | **22,453,829** | **20,493,851** | **17,862,489** | **19,981,092** |
| | **lost Rev.** | **0** | **3,472** | **17,859** | **17,850** | **37,811** | **66,993** | **60,532** | **95,935** | **84,344** | **106,545** | **90,163** | **110,767** | **143,952** | **158,787** | **226,819** | **189,302** | **142,971** | **242,344** | **191,414** | **177,813** | **251,333** | **133,892** | **200,879** | **160,351** | **229,766** | **356,751** | **377,433** | **341,759** | **299,993** | **316,847** |

Note: these data omitililary and non-grazed public lands (e.g. National Parks) so yield gap value may be slightly lower than the values presented in main text.
